# Activation of a Protein Kinase Via Asymmetric Allosteric Coupling of Structurally Conserved Signaling Modules

**DOI:** 10.1101/611772

**Authors:** Yuxin Hao, Jeneffer England, Luca Belluci, Emanuele Paci, H. Courtney Hodges, Susan S. Taylor, Rodrigo A. Maillard

## Abstract

Cyclic nucleotide binding (CNB) domains are universally conserved signaling modules that regulate the activities of diverse protein functions. Yet, the structural and dynamic features that enable the cyclic nucleotide binding signal to allosterically regulate other functional domains remain unknown. We use force spectroscopy and molecular dynamics to monitor in real time the pathways of signals transduced by cAMP binding in protein kinase A (PKA). Despite being structurally conserved, we find that the response of the folding energy landscape to cAMP is domain-specific, resulting in unique but mutually coordinated regulatory tasks: one CNB domain initiates cAMP binding and cooperativity, while the other triggers inter-domain interactions that lock the active conformation. Moreover, we identify a new cAMP-responsive switch, whose stability and conformation depends on cAMP occupancy. Through mutagenesis and nucleotide analogs we show that this dynamic switch serves as a signaling hub, a previously unidentified role that amplifies the cAMP binding signal during the allosteric activation of PKA.

## Introduction

Throughout evolution, nature has utilized conserved protein domains as regulatory signaling modules^1–6^. In multi-domain assemblies, these signaling modules communicate and transduce ligand binding signals to other functional domains, thereby enabling diverse responses to intracellular signaling cascades^7,8^. Cyclic nucleotide binding (CNB) domains are ubiquitous and structurally conserved signaling modules that regulate the activities of protein kinases, guanine nucleotide exchange factors, ion channels, and transcription factors in response to cyclic nucleotides^1^. To date, a general understanding of how the activity of CNB domains can be adapted to regulate a diverse array of protein functions remains rudimentary.

Here, we used optical tweezers in combination with steered molecular dynamics to study the mechanisms that link ligand binding and inter-domain communication with allosteric regulation of cAMP-dependent protein kinase A (PKA). PKA is an archetype of cyclic nucleotide-dependent protein kinases that is composed of regulatory and catalytic subunits (**Fig. 1a**)^9^. The phosphorylating activity of the catalytic subunit is allosterically driven by two CNB domains of the regulatory subunit, termed CNB-A and CNB-B^10–12^. cAMP binding starts in the CNB-B domain and enables binding of a second cAMP molecule to the CNB-A domain, resulting in a profound conformational change that unleashes the activity of catalytic subunits^10^.

**Figure 1.**
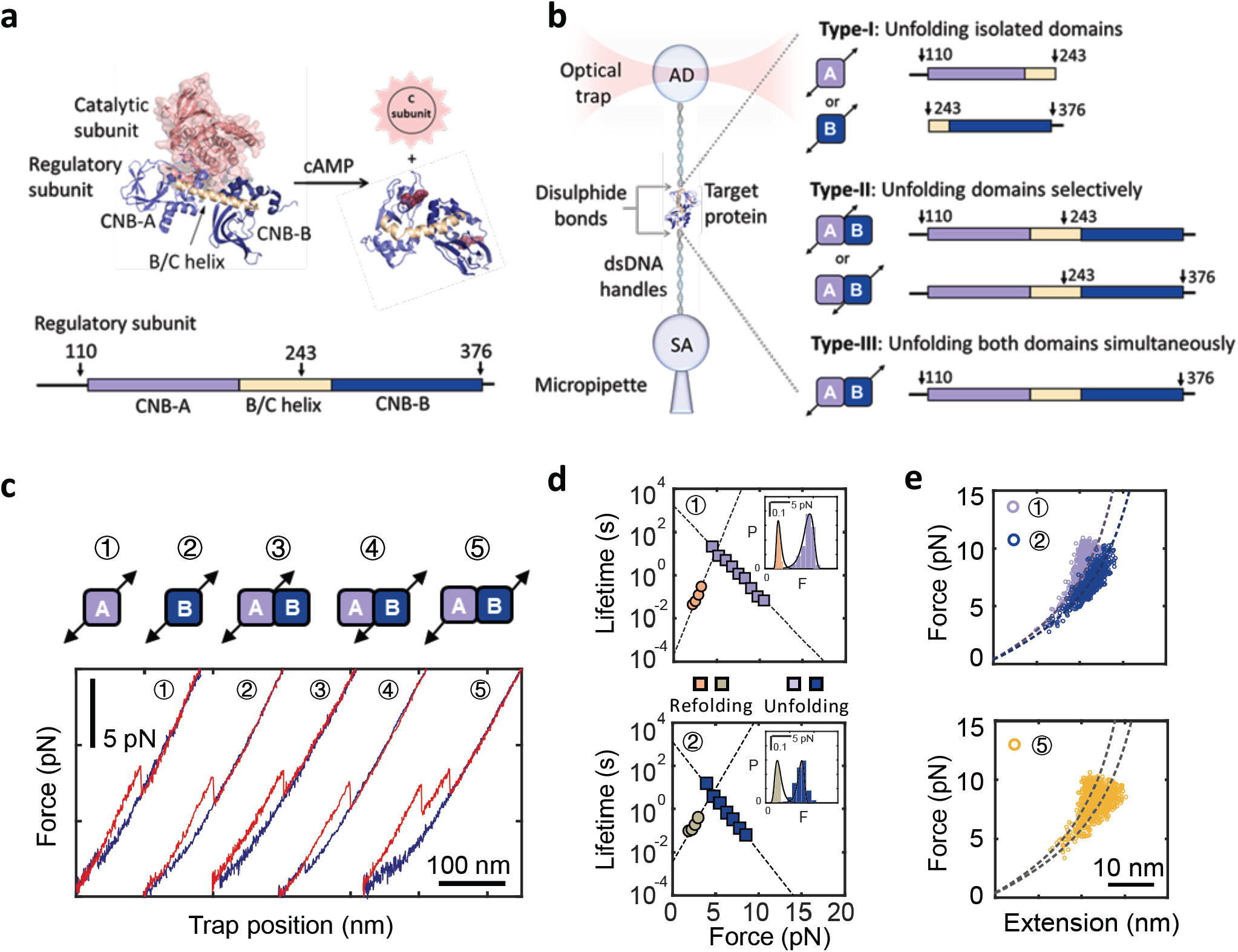
Experimental design to study allosteric activation in PKA. **a**, Structure of the inactive PKA holoenzyme (left)^8^ and active cAMP-bound regulatory subunit (right)^9^. The arrows on regulatory subunit domain organization indicate the residue positions for DNA handle attachment (bottom, arrows). **b**, Schematic representation of optical tweezers assay (left) and protein constructs used in this study (right). **c**, Force-extension curves for all pulling geometries in the apo state (unfolding in red; refolding in blue). **d**, Force-dependent folded state lifetimes and unfolding force probability distribution (inset) in the apo state. Black lines in insets are the unfolding force distribution reconstructed from force-dependent lifetimes. **e**, WLC analysis of changes in extension vs. force for the isolated CNB domains (top) and for the 1^st^ and 2^nd^ unfolding rips from the type-III construct (bottom). Dashed lines are the WLC curves for the CNB-A (purple) and CNB-B domains (blue). Numbering in 1d and 1e is the same as in panel 1c.

Our studies show that cAMP binding to the two CNB domains of PKA propagates a reorganization of inter-domain contact nodes that reshape the folding energy landscape of the protein. Changes in the energy landscape are unique to each CNB domain and arise from both ligand-binding and inter-domain interactions. We identify a division of labor among CNB domains: the CNB-B domain is responsible for initiating and triggering cAMP binding cooperativity while the CNB-A domain induces strong inter-domain interactions that lock the entire protein complex into its active conformation. Moreover, we identify a new cAMP-responsive structural element, the N3A motif, that switches in stability and conformation depending on cAMP occupancy and inter-domain contacts. Through mutagenesis and the use of cyclic nucleotide analogs, we show that this ligand-responsive switch is selective to cAMP and serves as a signaling hub, amplifying the cAMP binding signal during the allosteric activation of PKA. Altogether, this study illustrates how each structurally conserved CNB domain has evolved to carry out unique but mutually coordinated regulatory tasks in a macromolecular assembly. Our work reveals new operating principles for ligand-directed protein allostery mediated by widely conserved signaling modules.

## Results

### Multiple Pulling Geometries with Optical Tweezers Allow the Dissection of Folding Energy Landscape Parameters

To study the CNB domain communication mechanisms triggered by cAMP, we began by perturbing the free energy landscape of the PKA regulatory subunit with optical tweezers (**Fig. 1b, left**). We attached DNA handles via thiol chemistry to two cysteines engineered at specific positions in the protein (see Methods)^13,14^. The handle position determines the direction and region of the protein subjected to the force applied through the optical tweezers (i.e., pulling geometry)^15– 17^. We generated three PKA regulatory subunit constructs with unique pulling geometries to probe cAMP binding coupled to inter-domain interactions (**Fig 1b, right**). In type-I constructs, force is applied to the isolated CNB domains to study the effect of cAMP binding on the free energy landscape of each domain. In type-II constructs, force is applied selectively to one CNB domain in the presence of the neighboring one. This pulling geometry allows us to directly assess how cAMP binding induces inter-domain interactions, a strategy that would otherwise be inaccessible with bulk methods or single molecule fluorescence techniques. In type-III constructs, force is applied across both CNB domains simultaneously, allowing non-contiguous regions of the protein to respond to force, thereby probing long-range allosteric interactions, either in the presence or absence of cAMP.

We separately tethered each type of protein construct between two polystyrene beads in the optical tweezers (**Fig. 1b, left** and **Methods**). By gradually increasing and decreasing the tension across a single protein (“force ramp” pulling), we observed one or more rips in the resulting force-extension curves that correspond to unfolding and refolding events, respectively (**Fig. 1c**). In the apo state, the isolated CNB domains unfold at a similar average force, F_avg_ ∼ 7-9 pN, and with similar unfolding kinetic parameters: the lifetime of the folded state extrapolated to zero force, τ_0,F_, is 1.1-1.6×10^3^ s and the distance to the transition state, 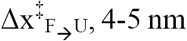, 4-5 nm (**Fig. 1d** and **Supplementary Table 1**)^18,19^. Analysis of refolding transitions show small differences, wherein the isolated CNB-A domain has a shorter unfolded state lifetime, τ_0,U_, and a longer 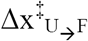 compared to the isolated CNB-B domain (**Supplementary Table 1**). The selective (type-II constructs) or simultaneous (type-III construct) mechanical manipulation of the CNB domains in the apo state showed indistinguishable unfolding and refolding kinetic parameters compared to their isolated counterparts (**Fig. 1e** and **Supplementary Fig. 1 and 2**), indicating that inter-domain interactions within the PKA regulatory subunit are negligible in the absence of cAMP^20^.

### Asymmetric Domain Stabilization Effects Triggered by cAMP Binding

In contrast to the results obtained in the apo state, the presence of cAMP revealed important differences between the two CNB domains. The unfolding force of the isolated CNB-B domain increases to F_avg_ = 12.0 ± 1.0 pN (**Fig. 2a, b**, N = 648), resulting in a ∼ 30-fold increase of τ_0,F_ (**Fig. 2c** and **Supplementary Table 2**). For the isolated CNB-A domain, F_avg_ = 17.4 ± 2.0 pN (**Fig. 2d, e**, N = 785) and τ_0,F_ increases by a factor of ∼ 7 (**Fig. 2f**). The kinetic stabilization conferred by cAMP is also observed during the refolding reaction; both CNB domains had a ∼ 4-fold decrease in τ_0,U_ (**Supplementary Fig. 1**). These effects illustrate that the minor structural differences between the two CNB domains (*r.m.s.d.* = 1.2 Å between Cα atoms) do not reflect the important differences in the folding energy landscape response to cAMP binding.

**Figure 2.**
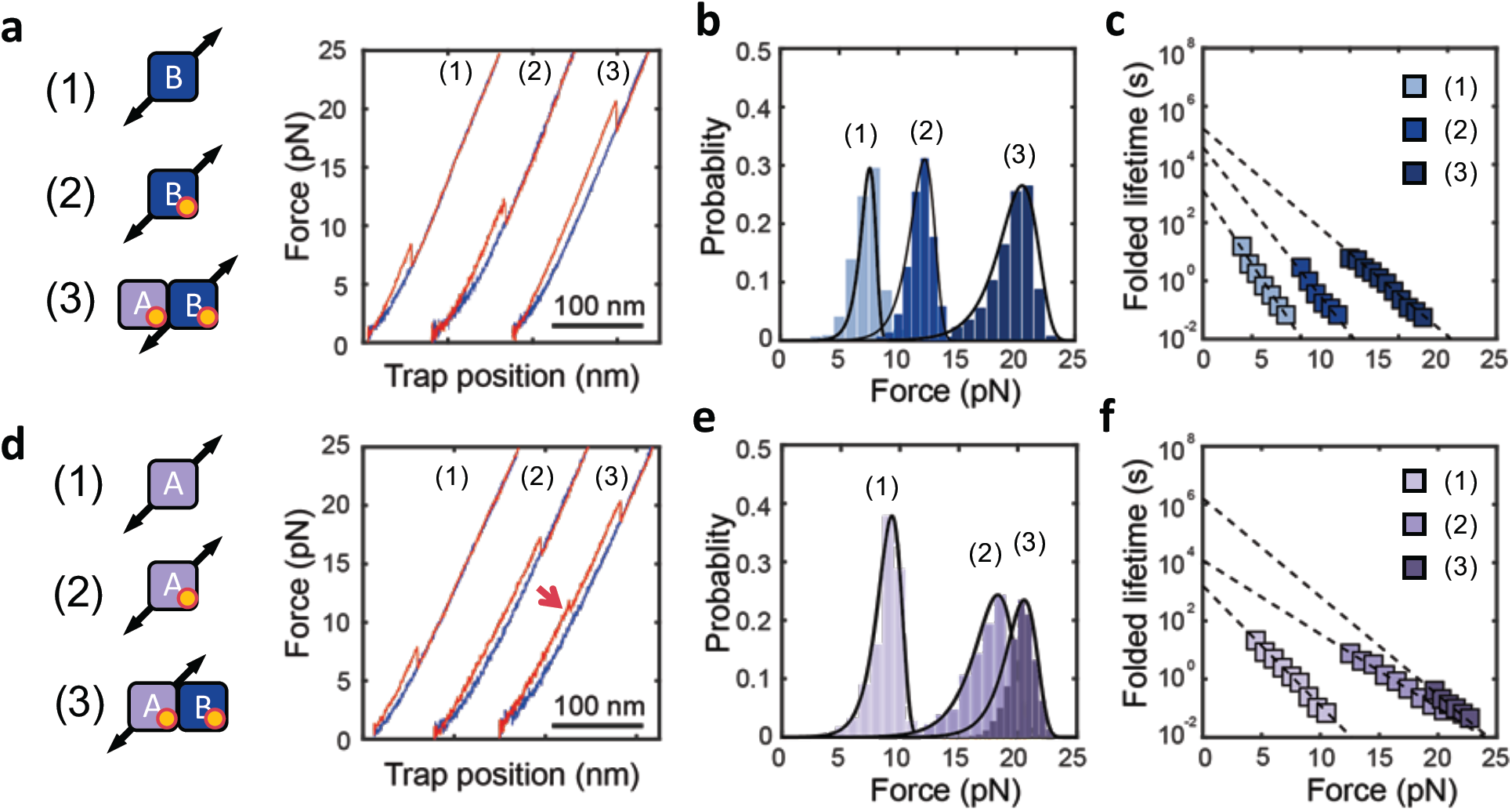
Selective allosteric effects initiated by cAMP binding. Force-extension curves **(a, d)**, unfolding force probability distributions **(b, e)**, and force-dependent folded state lifetimes **(c, f)** for the CNB-B (top) and CNB-A (bottom) domains. Numbering corresponds to the isolated CNB domains in the apo (1) or cAMP-bound states (2), and selective unfolding of the CNB domains bound to cAMP (3). The red arrow in panel 2d indicates the unfolding of the N3A motif.

### cAMP Induces Inter-Domain Interactions that Reshape the Folding Energy Landscape

Having characterized each isolated CNB domain, we studied inter-domain interactions triggered by cAMP using type-II constructs. We find that both CNB domains were stabilized by the presence of their counterpart when bound to the cyclic nucleotide (**Fig. 2**). Interestingly, the magnitude of stabilization was asymmetric (**Supplementary Table 2**): The CNB-A domain stabilizes the CNB-B domain by an additional ∼ 8 pN, resulting in F_avg_ = 19.7 ± 1.6 pN (N = 1518) and a 4-fold increase in τ_0,F_. The presence of the CNB-B domain induces a mechanical stabilization to the CNB-A domain of ∼ 3 pN, resulting in F_avg_ = 20.3 ± 1.4 pN (N = 1152) and a 160-fold increase in τ_0_. In the refolding reaction, the presence of the neighboring domain decreases τ_0,U_ by 150-fold and 10-fold to the CNB-B and CNB-A domains, respectively (**Supplementary Fig. 1**). These results show that cAMP binding induces specific but coordinated effects, wherein the CNB-B domain stabilizes the folded state of the CNB-A domain, and the CNB-A domain destabilizes the unfolded state of the CNB-B domain. Therefore, the cAMP-dependent communications between the CNB domains is bidirectional and asymmetric, highlighting a unique role for each domain in the activation mechanism of PKA.

### Identification of a cAMP-Responsive Dynamic Switch

We hypothesized that changes in contour length upon unfolding (ΔL_c_) might also reveal important differences in the native folded structures of the CNB domains upon binding cAMP. While the mechanical unfolding of the CNB-B domain in all three types of constructs had a ΔL_c_ of ∼ 50 nm, corresponding to a fully folded domain (**Supplementary Table 2**), the CNB-A domain displayed a more complex behavior. The isolated CNB-A domain in the apo state had the expected ΔL_c_ of 45 nm based on the crystal structure^10^. However, the value of ΔL_c_ decreased to 30 ± 3 nm in the presence of cAMP, indicating that a region of the domain was destabilized upon ligand binding (**Supplementary Table 2**).

We sought to identify which region or secondary structures of the CNB-A domain become unstable upon cAMP binding. The structure of the CNB-A domain is composed of a β-sandwich fold that forms the cAMP binding pocket, and three N-terminal α-helices termed N3A motif.^10,11^ The N3A motif contains ∼ 30 amino acids, which matches the amount of polypeptide that became unstable after cAMP binding. To test whether the N3A motif is destabilized by cAMP binding, we used two distinct type-III constructs, one with DNA handles attached at residue positions flanking both CNB domains entirely (S110C/S376C) and another construct with handles flanking both CNB domains except the N3A motif (D149C/S376C). The two constructs displayed two major unfolding rips corresponding to the CNB domains, but only the unfolding trajectory of the S110C/S376C construct revealed a small, reversible transition at ∼ 11 pN with a ΔLc of 13 nm (**Fig. 3a**). The lack of such small transition in the D149C/S376C construct, which does not directly probe the N3A motif, provides evidence that the secondary structures in the CNB-A domain that become unstable upon cAMP binding correspond to the N3A motif (**Supplementary Fig. 3**). Steered molecular dynamics (SMD) starting from the X-ray structure corroborate our experimental observations (**Methods**). The SMD trajectories show that the N3A motif unravels first while the rest of the CNB domains remained stably folded in their original cAMP-bound conformation (**Fig. 3b-right, and Supplementary Movie 1**). In contrast, SMD trajectories in the absence of cAMP several interdomain interactions were lost (**Supplementary Fig. 4**), resulting in the detachment of the two CNB domains before any secondary structure unfolds, including the N3A motif (**Fig. 3b-left, and Supplementary Movie 2**).

**Figure 3.**
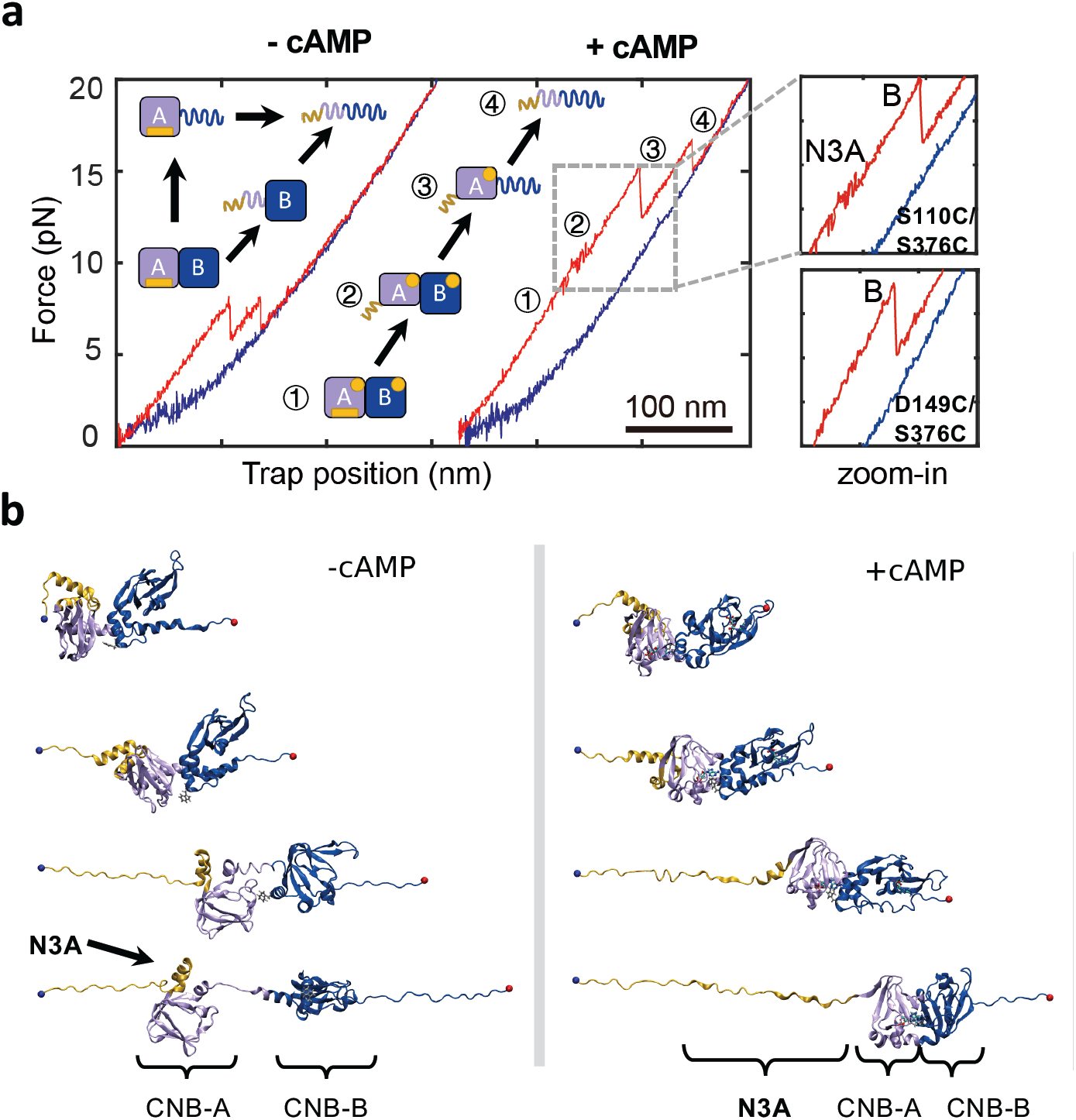
Dual unfolding pathways of PKA regulatory subunit depending on cAMP occupancy. **a**, Force-extension curve for the cAMP-free and cAMP-bound regulatory subunit (type-III construct). The cartoon represents the structural transitions occurring during the unfolding trajectory. Zoomed in are the unfolding trajectories of type-III constructs S110C/S376C and D149C/S376C. A detailed analysis of the forces for each unfolding transition refer to Supplementary Information and Supplementary Table 4. **b**, SMD simulation snapshots of the cAMP-bound (top) and apo (bottom) regulatory subunit. Yellow: N3A motif; Purple: CNB-A; Dark blue: CNB-B domain.

### Conformational Switching of the N3A Motif Requires Integrity of Inter-Domain Contacts

In contrast to the results obtained with the isolated cAMP-bound CNB-A domain, the type-III S110C/S376C construct show that the N3A motif is properly folded in the context of the entire regularly subunit. Moreover, a close inspection of trajectories obtained with the selective manipulation of the cAMP-bound CNB-A domain (type-II construct) revealed a two-step unfolding process instead of a single rip (**Fig 2d, red arrow**). The additional rip had a ΔL_c_ of ∼ 13 nm, similar to that of the N3A motif. These observations indicate that the CNB-B domain enables the refolding of the N3A motif in the presence of cAMP.

Because our optical tweezers assay permitted to control the sequence of events in the unfolding reaction, we used the type-III S110C/S376C construct to determine whether the N3A motif can refold while the CNB-B domain remains in the unfolded state (**Fig. 4a**). In this experiment, we applied a force up to 15 pN to unfold both the N3A motif and the CNB-B domain, but not the CNB-A domain. The force was then decreased to 5 pN to maintain the CNB-B domain in the unfolded state (refolding transitions begin at forces < 2 pN). After ten or more pulling and relaxation cycles between 5-15 pN, we did not observe any small, reversible transitions at ∼ 11 pN, which would have corresponded to a folded N3A motif while the CNB-B domain remained unfolded. Thus, we find that the CNB-B domain is strictly required for the N3A motif to refold. This result is in agreement with the structure of the cAMP-bound regulatory subunit that shows the N3A motif docks into a cleft formed between the CNB domains,^11^ establishing several surface contacts not only with the CNB-A domain and the BC helix,^21^ but also with the CNB-B domain (**Fig. 4b**).

**Figure 4.**
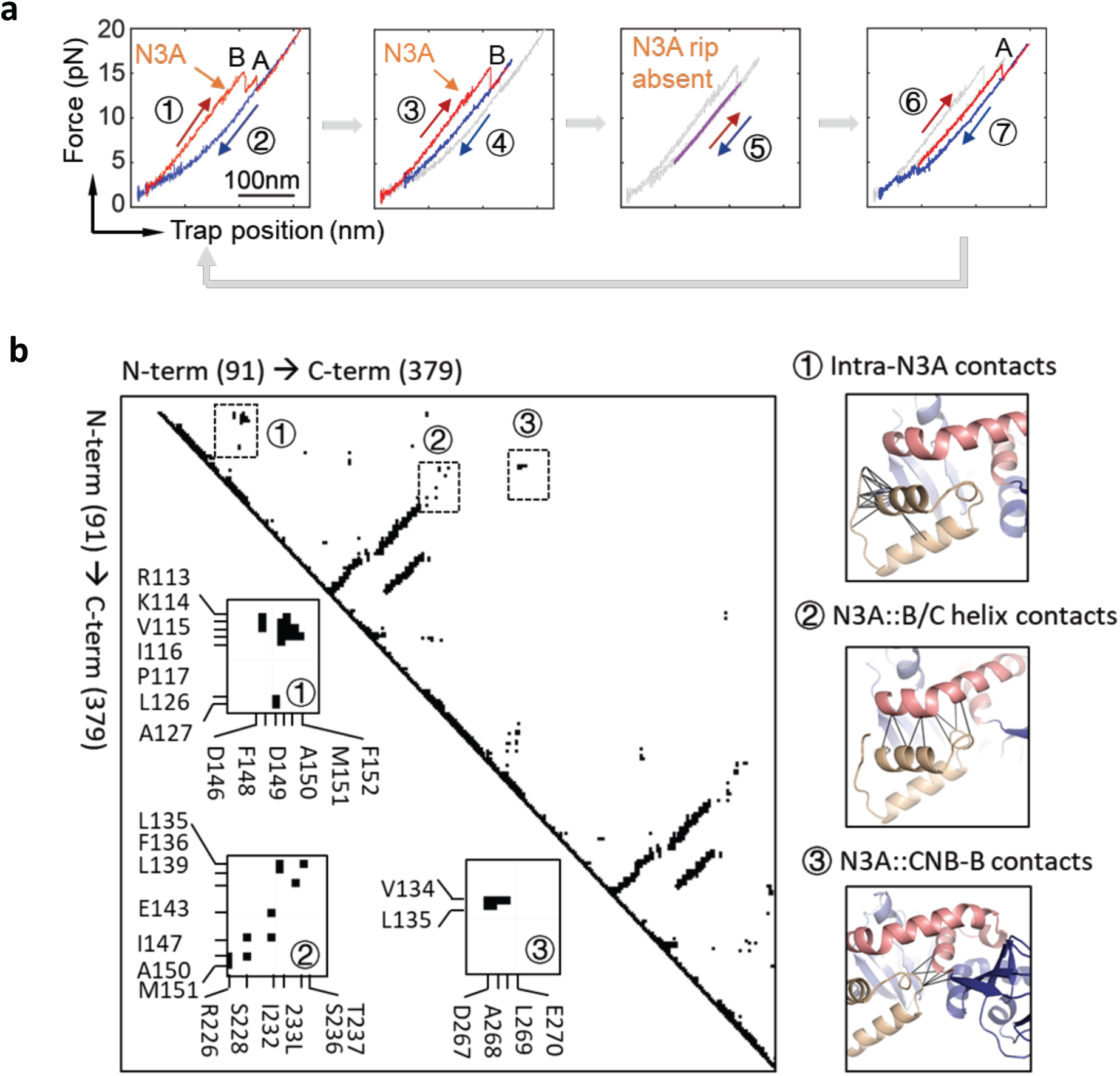
cAMP-bound CNB-B is required for N3A motif to refold. **a**, 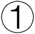 The regulatory subunit unfolds (red) and 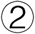 refolds (blue), revealing the 1^st^ reversible transition corresponding to the N3A motif (orange arrow). 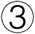 In the following cycle, the regulatory subunit was stretched until the N3A motif and the CNB-B domain unfold, while the CNB-A domain remains folded. 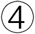 The force was decreased to 4 pN, a force that does not allow the CNB-B domain to refold. 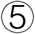 The force was then oscillated between 6-15 pN for several cycles (∼ 20) to test whether the N3A motif was able to refold while the CNB-B domains remains unfolded. 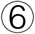 The force was increased to 20 pN to unfold the CNB-A domain. 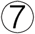 The force was decreased to 1 pN, allowing the complete protein to refold, and begin another set of experiments. The trajectories in gray represent the unfolding pathways from the immediately previous cycle, thereby serving as reference on the progression of experiment. **b**, Pairwise contact map comparing the interaction established by the N3A motif in the regulatory subunit of PKA (left). The contacts established by the N3A motif were obtained using a 8 Å cutoff (right): 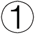 contacts established by residues within the N3A motif; 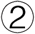 contacts between the N3A motif and the B/C helix; and 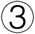 contacts between the N3A motif and the CNB-B domain. Cartoons rendering the three sets of contacts are shown next to the contact map.

### Intermediate cAMP-Bound States Reveal Stepwise Stabilization Mechanism Between CNB Domains

Since cAMP binds to PKA in a sequential fashion^10^, thereby populating intermediate-cAMP-bound species, we investigated the coupling between the folding status of the N3A motif and inter-domain interactions in conditions where only one CNB domain is bound to cAMP. To obtain force-extension curves of intermediate cAMP-bound states, we used the type-III construct S110C/S376C and titrated cAMP between 1-150 nM (**Fig. 5a**). We find that the CNB-B domain bound to cAMP increases both F_avg_ by ∼ 1 pN (KS Test, *p* ≅ 0) and τ_0,F_ by 2-fold to the cAMP-free CNB-A domain (**Fig. 5b, top**). The CNB-A domain bound to cAMP induces a larger stabilization to the cAMP-free CNB-B domain, increasing F_avg_ by ∼ 3 pN and τ_0,F_ by 3-fold (**Fig. 5b, bottom**). These results show that cAMP binding to one CNB domain is sufficient to initiate stabilizing inter-domain interactions with the neighboring apo CNB domain, but compared to the fully cAMP-bound state these interactions are partial in magnitude (**Supplementary Table 3**). Moreover, we find that these partial inter-domain interactions are insufficient to drive the folding of the N3A motif between the two CNB domains, i.e., analysis of ΔL_c_ using the WLC model shows that the cAMP-bound CNB-A domain interacting with the cAMP-free CNB-B domain does not have a folded N3A motif (**Fig. 5c, top**). A similar analysis revealed that cAMP binding to the CNB-B domain does not elicit unfolding of the N3A motif in the cAMP-free CNB-A domain (**Fig. 5c, bottom**). These results strongly support our previous observations showing that unfolding of the N3A motif is solely coupled to cAMP binding to the CNB-A domain, and that the following refolding step of the N3A motif requires the presence of the cAMP-bound CNB-B domain (**Fig. 4a**).

**Figure 5.**
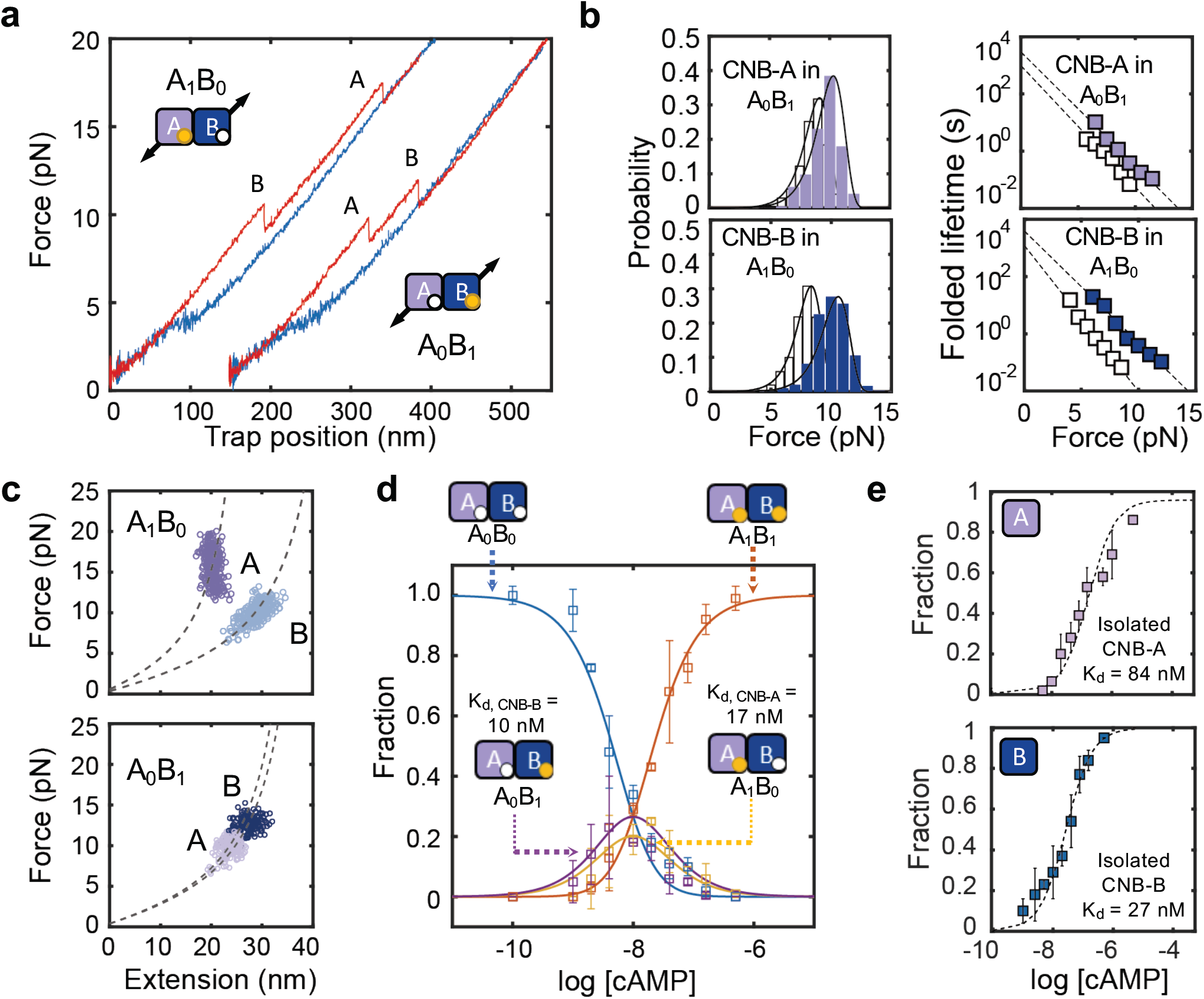
Stepwise Stabilization of CNB Domains in Partial cAMP-bound States. **a**, Force-extension curves of intermediate-liganded states (A_1_B_0_ and A_0_B_1_) for the type-III regulatory subunit S110C/S376C. **b**, Unfolding force probability distribution and force-dependent folded state lifetimes for intermediate-liganded states: CNB-A domain in A_0_B_1_ (top) and CNB-B domain in A_1_B_0_ (bottom). The corresponding isolated domains in the apo state (white bars and symbols) are shown for comparison. Solid lines are the unfolding force distribution reconstructed from force-dependent lifetimes. **c**, WLC analysis of changes in extension upon unfolding vs. force in A_1_B_0_ and A_0_B_1_ for the CNB-A and CNB-B domains (dashed lines). **d**, cAMP titration plot showing the fraction of apo (A_0_B_0_), intermediate (A_1_B_0_, A_0_B_1_) and fully bound (A_1_B_1_) species. Lines correspond to the global fit to the equations for each species. **e**, Fractional titration plot of isolated CNB-A (top) and CNB-B (bottom) domains (type-I constructs). The error bar corresponds to the standard deviation of 5-10 different molecules.

We also find that partial inter-domain interactions initiated by on-pathway intermediate cAMP-bound states have important functional consequences in terms of cAMP binding affinities and cooperativity. By directly counting unfolding trajectories corresponding to the apo, intermediate and fully bound species as a function of cAMP concentration (**Supplementary Fig. 5**), we built a single-molecule titration curve, globally fitted the equations for each population species, and determine the microscopic binding constants and cooperativity parameter (**Fig. 5d** and **Supplementary Info**). For the first cAMP molecule, the CNB-B domain has a dissociation constant (K_d,CNB-B_) of 10 ± 1 nM and the CNB-A domain has a K_d,CNB-A_ = 17 ± 1 nM. The affinity for the second cAMP molecule for either CNB domain increases ∼ 3-fold, indicating positive binding cooperativity. Importantly, the K_d_ values of the CNB domains are 3 and 5 times lower than those corresponding to the isolated domains (see **Supplementary Info**), indicating that as part of the regulatory subunit the CNB domains bind cAMP more tightly (**Fig. 5e**).

### Conformational Stability and Dynamics of the N3A Motif Are Critical During the Allosteric Activation of PKA

Our results portray the N3A motif as a ligand-responsive molecular switch that toggles between different conformations depending on cAMP occupancy and specific domain contacts. This unique character led us to hypothesize that the N3A motif is a critical structural element that mediates cAMP-dependent cooperative interactions between the CNB domains. We tested this hypothesis by placing the mutation R241A in the B/C helix that connects both CNB domains. In the wild type structure bound to cAMP, R241 interacts with D267 in the CNB-B domain and E200 in the CNB-A domain, thereby bringing the two CNB domains into close proximity for the N3A motif to dock (**Fig. 6a, left**)^11,22,23^. In the absence of cAMP, unfolding trajectories of R241A using a type-III construct (S110C/S376C) show indistinguishable unfolding parameters compared to wild type (**Supplementary Fig. 6**). In the presence of cAMP, however, the trajectories of R241A revealed an unfolding pathway that looked similar to that of wild type (**Fig. 6a, right** and **Supplementary Fig. 6**), despite some important quantitative differences. Specifically, the average unfolding force for the CNB-B domain in the mutant protein was ∼ 2.5 pN lower compared to wild type, which results in a 3-fold reduction of τ_0,F_ (**Fig. 6b**). These values are indistinguishable from those obtained with the isolated CNB-B domain bound to cAMP (**Supplementary Table 4**), indicating that R241A largely eliminates inter-domain interactions initiated by the cyclic nucleotide. SMD trajectories also show that the first event in the unfolding pathway of R241A is the detachment of the cAMP-bound CNB domains, instead of the unraveling of the N3A motif (**Fig. 6c, Supplementary Fig. 4** and **Supplementary Movie 3**).

**Figure 6.**
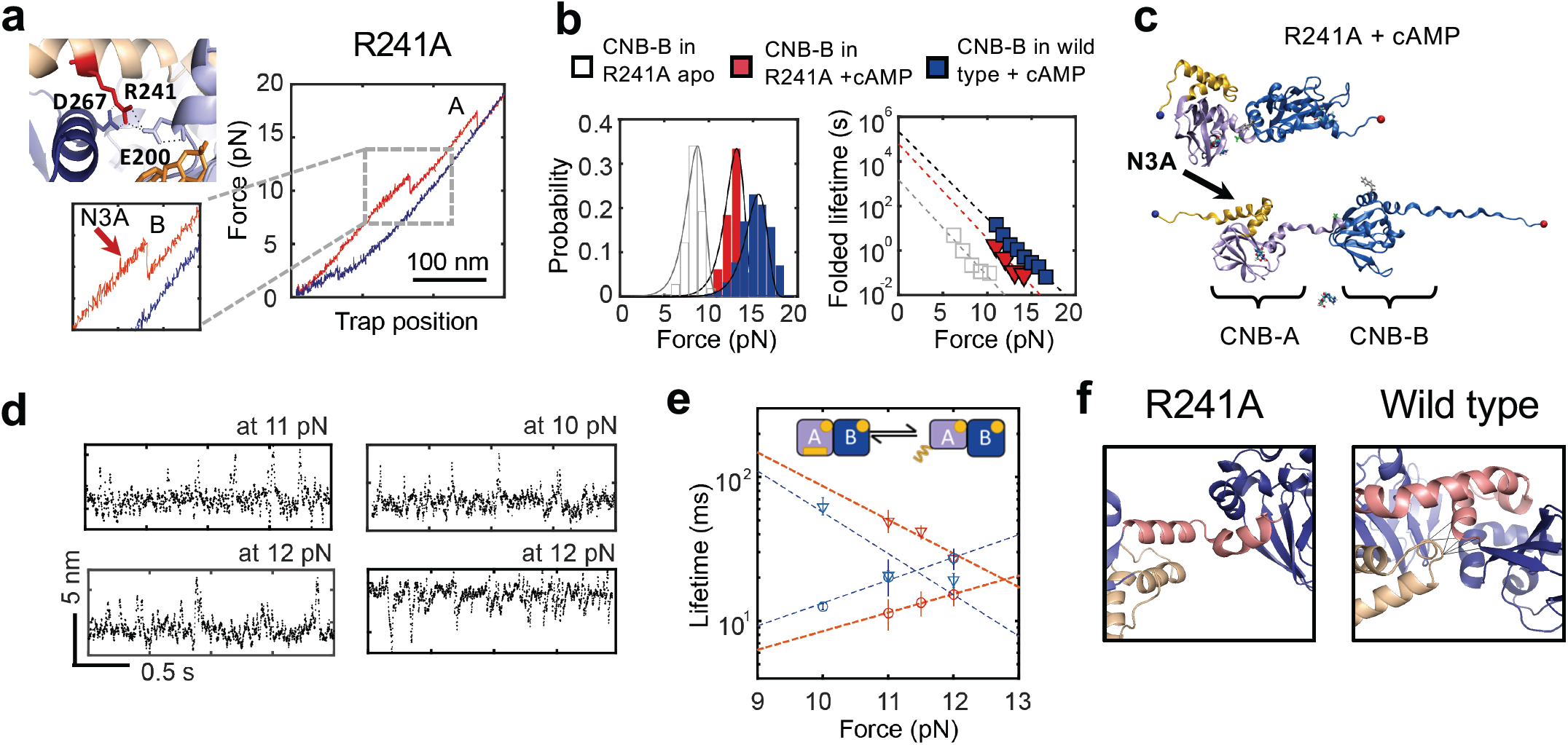
Perturbation of allosteric networks in PKA by Mutation R241A. **a**, Residue R241 interacts with both CNB domains through E200 and D267 (PDB 1RGS). Force-extension curve for R241A bound to cAMP (type-III construct S110C/S376C). Zoomed-in is the unfolding rip corresponding to the N3A motif. **b**, Unfolding probability distributions and force-dependent folded state lifetimes for the CNB-B domain in R241A apo (gray) or bound to cAMP (red). For reference the wild type data was included (blue). Solid lines are the unfolding force distribution reconstructed from force-dependent lifetimes. **c**, SMD simulation snapshot of the cAMP-bound R241A protein. **d**, Representative force-clamp trajectories of the N3A motif in R241A (left) and wild type (right). **e**, Force-dependent lifetimes of the N3A motif in the folded (triangles) and unfolded (circles) states for R241A (red) and wild type (blue). **f**, The mutation R241A (left) hinders interactions established with the CNB-B domain seen in wild type (right).

To further dissect the role of folding dynamics and conformation of the N3A motif in inter-domain interactions, we conducted “force-clamp” experiments, wherein the protein was held at varying constant forces between 10-12 pN, and changes in extension due to unfolding and refolding of the N3A motif were monitored as a function of time (**Fig. 6d**). Analysis of these trajectories using a two-state Bayesian Hidden Markov model (BHMM)^24,25^ revealed that the folded-state lifetime was ∼2 times longer and the unfolded-state lifetime was ∼2-fold shorter for R241A, indicating that the N3A motif in the mutant protein is less dynamic than in wild type (**Fig. 6e**). In addition, the accompanying ΔL_c_ between the folded and unfolded states for R241A was 6.5 ± 1.1 nm and for wild type was 9.5 ± 0.5 nm. The difference in folding dynamics and ΔL_c_ indicates that the N3A motif in R241A is stably folded. But because the mutation eliminates cAMP-dependent inter-domain interactions (**Fig. 6b**), it is likely that the folded N3A motif is not docked between the two CNB domains (**Fig. 6f**), thereby impeding the PKA regulatory subunit to attain its final cAMP-bound conformation. Thus, our single-molecule studies show that R241A mutant imparts cAMP-dependent functional deficiencies^26^ due to a disruption of the conformational dynamics of the N3A motif and its ability to serve as an efficient cAMP-responsive molecular switch.

### Dynamic Switching of the N3A Motif is Selective to cAMP

To study the contribution of the N3A motif conformational switching mechanism towards cyclic nucleotide selectivity, we mechanically manipulated the CNB domains individually (type-I constructs) or simultaneously (type-III construct) in the presence of cGMP. Both CNB domains bound to cGMP show unfolding parameters (F_avg_, τ_0,F_, and 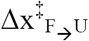) that lie in between the values obtained with and without cAMP, indicating partial intra-and inter-domain stabilization effects (**Fig. 7a, b** and **Supplementary Table 5**). Interestingly, the isolated cGMP-bound CNB-A domain had a greater ΔLc than its cAMP-bound counterpart (37 nm and 30 nm, respectively), indicating that the N3A motif is not negatively coupled to cGMP, but instead unfolds as a single cooperative unit together with the rest of the domain. In agreement with this interpretation, the force-extension curves using a type-III construct with cGMP do not show the small, reversible transition characteristic of the N3A motif (**Fig. 7a**). Rather, the trajectory revealed two unfolding rips with ΔL_c_ values that reflect the mechanical denaturation of the full-length protein (**Fig. 7c** and **Supplementary Fig. 7**). These results provide direct experimental evidence that nucleotide selectivity not only involves previously described defects in binding affinity and cooperativity^27^, but also an attenuation of inter-domain interactions and decoupling of cyclic nucleotide binding from the conformational switching of the N3A motif.

**Figure 7.**
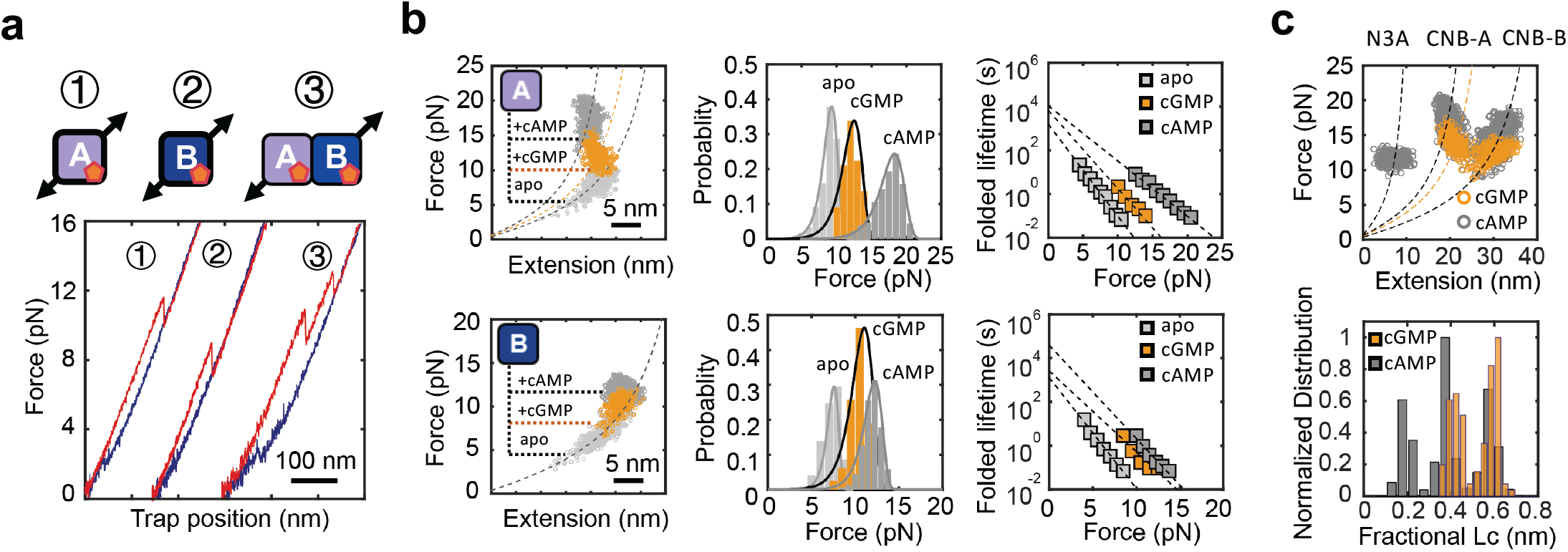
Perturbation of allosteric networks in PKA by cGMP. **a**, Force-extension curves of cGMP-bound protein constructs. **b**, WLC analysis of changes in extension upon unfolding vs. force (left), unfolding force probability distributions (center) and force-dependent folded state lifetimes (right) for the isolated CNB-A (top) and CNB-B (bottom) domains in the apo (light gray), cGMP-bound (orange), and cAMP-bound (dark gray) states. Solid lines in center panels are the unfolding force distribution reconstructed from force-dependent lifetimes. **c**, WLC analysis of changes in extension upon unfolding vs. force (top) and fractional contour length (bottom) of the regulatory subunit bound to cGMP (orange) and cAMP (gray). Dashed lines are the WLC curves for the N3A motif and the two CNB domains with cAMP (gray) and cGMP (orange).

## Discussion

The uncovered intra-and inter-domain communication network that is triggered by cAMP binding cannot be easily inferred from the crystal structure^12^. The network of interactions in PKA involves bidirectional communication that is asymmetric in magnitude, and includes both positive (stabilizing) and negative (destabilizing) coupling interactions that are fine-tuned to attain the final cAMP-bound conformation. Positive coupling promotes stabilizing interfacial interactions between CNB domains. cAMP binding stabilizes the CNB-B domain from 7.6 kcal/mol to 10.4 kcal/mol, and the presence of the neighboring cAMP-bound CNB-A domain provides another 3.2 kcal/mol. We estimate that cAMP binding stabilizes the CNB-A domain from 9.4 kcal/mol to 11.5 kcal/mol, and inter-domain interactions confer an additional 4.4 kcal/mol (**Fig. 8a** and **Supplementary Table 1,2**). Negative coupling triggered by cAMP binding effectively melts interactions established between the N3A motif and the catalytic subunit, thereby facilitating the dissociation of the PKA complex. Lastly, the N3A motif folds back between the cAMP-bound CNB domains with a stability of 5.5 kcal/mol. Within this interaction network, we identified the unique routes of domain communication that are disrupted either by R241A or cGMP (**Fig. 8b**). Because the N3A motif is found in many other cyclic nucleotide binding proteins^28^, the uncovered conformational switching mechanism may be a widespread strategy not only to ensure the completion of the allosteric activation process but also to provide ligand selectivity.

**Figure 8.**
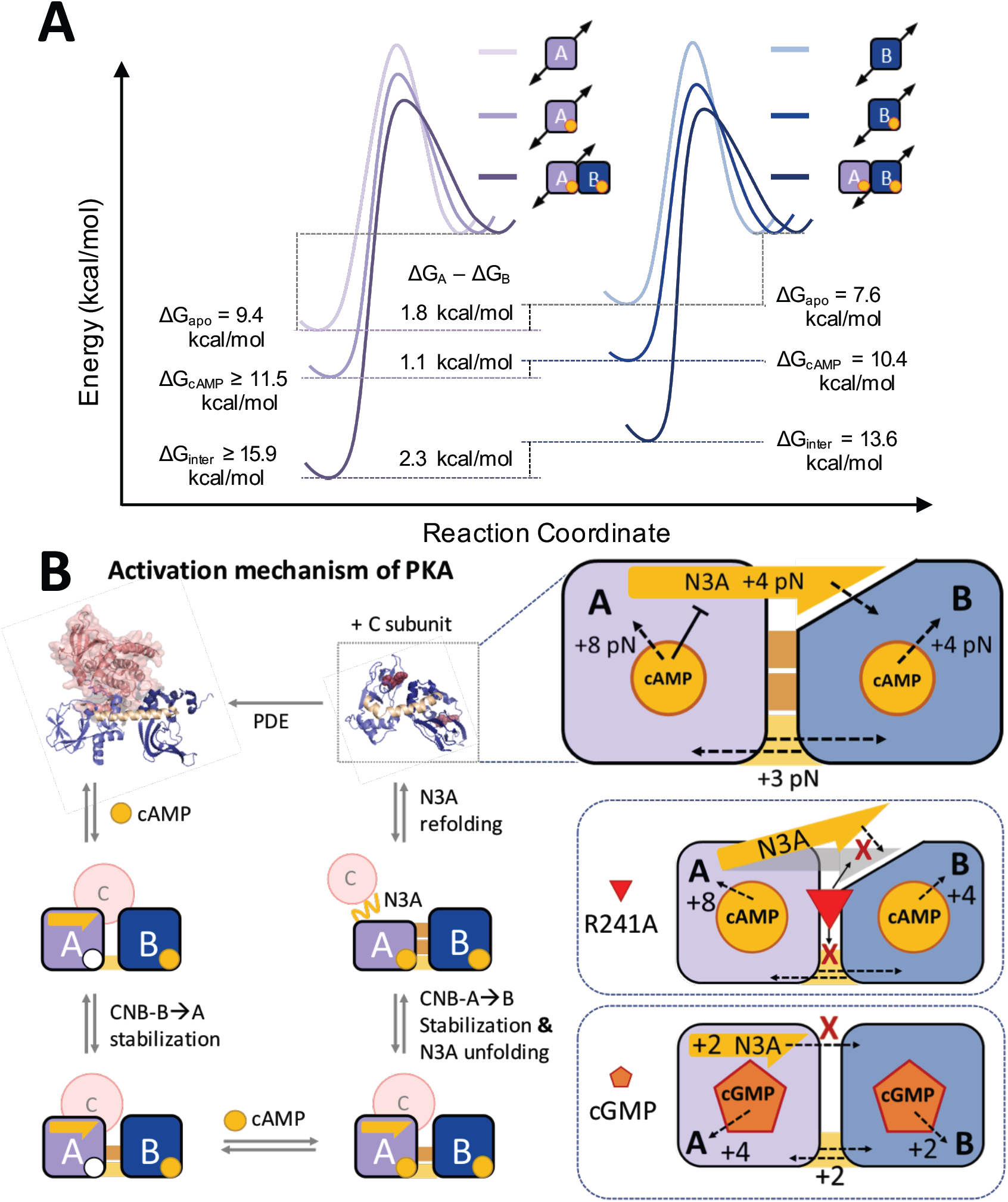
Activation of PKA through selective stabilization of CNB domains. **a**, Unique response of the energy landscape of the CNB domains due to cAMP binding and inter-domain contacts. The height of the energy barriers reflect the folded and unfolded state lifetimes of the CNB domains in the different configurations. **b**, Left: Activation mechanism of PKA showing intermediate states identified in this study. Right-top: Interaction network initiated by cAMP binding involves stabilizing (dashed black arrows) and destabilizing (flat black arrow) effects. Right-bottom: Interactions routes disrupted by R241A and cGMP.

In conclusion, our studies reveal a novel allosteric regulation mechanism, in which co-existing positive and negative coupling interactions initiated by cAMP binding are coordinated to sequentially dissociate the PKA complex, commencing from the CNB-B domain, which binds cAMP first and enables binding of the cyclic nucleotide to the CNB-A domain^10^, and finishing with the conformational switch of the N3A motif. Like unzipping “Velcro”, this mechanism efficiently peels off the two PKA subunits, where the strong inter-subunit interaction is the result of several smaller interactions that can be broken sequentially, hence the dissociation of the two subunits does not require the crossing of a large free energy barrier but of many small ones. We anticipate similar mechanisms in other protein kinases with catalytic subunits that require the dissociation of regulatory signaling modules^29^. Moreover, the allosteric mechanisms we describe here may be amplified in the PKA hetero-tetramer composed of two regulatory and two catalytic subunits, where the potential cross talk between PKA subunits is expanded. The single-molecule approach exploiting optical tweezers in conjunction of molecular dynamics simulations presented here can be extended to map allosteric effects of disease mutations or inhibitor binding in other kinases or multi-domain assemblies^30–34^.

## Online Methods

### 1. Mutagenesis, Expression and Purification of the PKA Regulatory Subunit and Isolated CNB Domains

The PRSET plasmid harboring the *Bos taurus* regulatory subunit gene, isoform RIα containing residues 91 to 379 of the full-length sequence was used. To obtain isolated CNB domains, the sequence of the neighboring CNB domain was deleted by site-directed mutagenesis (QuikChange II Agilent). The isolated CNB-A domain contains residues 91 to 243. The isolated CNB-B domain contains residues 243 to 379. The mutations C345A and C360A were introduced in the CNB-B domain to prevent undesired reactions with the thiol-modified dsDNA handles. To manipulate each individual CNB domain (Type-I constructs) we introduced the mutations S110C/M243C and M243C/S376C for the isolated CNB-A and CNB-B domains, respectively. To manipulate either the CNB-A domain or the CNB-B domain selectively (Type-II constructs) we introduced into the regulatory subunit the mutations S110C/M243C and M243C/S376C, respectively. To manipulate both CNB domains simultaneously (Type-III construct), we introduced into the regulatory subunit the mutations S110C/S376C. All the protein constructs were expressed in BL21(DE3) (NEB) and purified as described previously^11,20,35^. Briefly, the protein was expressed in BL21(DE3) competent cells overnight at 18°C with 1 mM IPTG. The cells were in 20 mM MES, 100 mM NaCl, 2 mM EGTA, 2 mM EDTA, 5 mM DTT, pH 6.5 and the spin supernatant was precipitated with 40% ammonium sulfate before binding to a cAMP-coupled agarose resin. The protein was eluted from the resin with cGMP and run on a size exclusion column to remove the excess cGMP.

### 2. Attachment of dsDNA Handles to Protein Constructs

We followed a protocol previously published by the Maillard laboratory^14^. Briefly, the purified target protein was concentrated to ∼ 5 mg/mL in 10 mM DTT to reduce all cysteine residues. The protein solution was run through 3 Micro Bio-Spin columns (Bio-Gel P6, Biorad) to remove the DTT before adding 10 mM 2,2’-dithiodipyriine (DTDP, Sigma) for 2 hours at room temperature. The unreacted DTDP was removed from the modified protein using 3 additional Micro Bio-Spin columns. The DTDP-activated protein and two different 30-bp 5’-Thiol-modified dsDNA oligos were combined in a 1-to-1 molar ratio and incubated overnight at 4°C. The resulting protein-oligo chimera was stored at -80° C. Each 30-bp 5’-Thiol-modified dsDNA oligo used in the formation of the protein-oligo chimera has a unique non-palindromic overhang that is used to ligate 350-bp dsDNA handles modified with either biotin or digoxigenin in their 5’ end. The cross-linking reaction was done in the DNA crosslinking buffer (50 mM Tris, 100 mM NaCl, pH 7.6). Single molecule optical tweezers measurements of protein constructs in the apo state were obtained by gradient elution (using cAMP) of the protein-oligo chimera from a cAMP-coupled agarose resin. The first eluted samples have an initial cAMP concentration of 20 μM, which is further diluted to a final concentration of ∼ 0.02 nM (100-fold < *K*_d_)^36^ before using it in the optical tweezers experiment.

### 3. Optical Tweezers Measurements

All data was collected in a MiniTweezers instrument^37^. Measurements were carried out in DNA crosslinking buffer. The protein sample with dsDNA handles was mixed with of 3.1 µm polystyrene beads (Spherotech) coated with anti-digoxigenin antibodies (termed AD bead) for 5 min at room temperature. The sample is diluted to 1 mL before applying it to the optical tweezers microchamber. The optical tweezer experiments were performed in DNA crosslinking buffer in a temperature-controlled room at 20 °C. A 2.1 μm bead coated with streptavidin (termed SA bead) is trapped on the tip of a micropipette, whereas the AD bead conjugated with the sample is trapped in the optical trap. To form a single tether, the AD bead in the optical trap was moved towards the SA beads on the micropipette tip. A single tether is confirmed by observing overstretching of the DNA handles at ∼ 65 pN^38^. Data was collected in two modes:

a. Force ramp experiments: To mechanically manipulate the target protein construct, we moved the AD bead in the optical trap away and towards the SA bead on the micropipette tip, which results in force-extension curves. The experiment was conducted at a constant pulling velocity of 75 nm/s, with a 10 second refolding time at 2 pN. Data was collected at a sampling rate of 200 Hz. Rupture forces representing the cooperative unfolding of one or more protein domains and their associated extension changes were analyzed using a custom-built program implemented in MATLAB software. Unfolding force probability distributions obtained from force-extension curves were transformed to folded state lifetimes as a function of force and analyzed using the methodology described by Dudko *et al*^18,19^. (Full details are provided in the Supplementary Information)
b. Force clamp experiments: the protein was stretched and then held at a desired constant force using constant-force feedback loop. Changes in position at a particular force were recorded at a frequency of 500 Hz for the R241A mutant and 1000 Hz for the wild type protein. The data from force clamp experiments was analyzed using a Bayesian Hidden Markov Model (BHMM) approach^24,39^.

### 4. Molecular Dynamics (MD) and Steered Molecular Dynamic (SMD) Simulation

MD and SMD simulation were performed with NAMD (v2.12)^40^. The CHARMM27 force field^41^ was used for the protein and counterions and the TIP3P^42^ for water. Parameters for cyclic adenosine monophosphate (cAMP) were obtained with the CHARMM general force field (CGenFF)^43^. The holo state was modeled starting from the X-ray structure (PDB code: 1RGS)^11^. The apo state was obtained by removing cAMP from both the binding sites. Regulatory subunit of protein kinase A (PKA) with and without cAMP were solvated in a cubic box of 90 Å side. Systems included 24,200 water molecules and counterions were added to guarantee charge neutrality. The bonds between hydrogens and heavy atoms were constrained with the SHAKE algorithm^44^. The r-RESPA multiple time step method^45^ was employed where long-range electrostatic interactions are updated every 4 fs, and all the other interactions every 2 fs. Periodic boundary conditions (PBC) were used and the long-range electrostatics was treated with the particle-mesh-Ewald (PME) method^46^ using a grid of 81×81×81. The cut off for non-bonded interactions was set to 10 Å, and the switching function was applied to smoot interactions between 9 and 10 Å. Simulations were conducted in the NPT ensemble. Temperature was set to 310 K through a Langevin thermostat^47^ and pressure was set to 1 atm through a isotropic Langevin piston manostat^48^. The systems were first minimized (2000 steps of conjugate gradient) and equilibrated for 800 ps with the atoms of protein restrained to their initial positions. Production runs for both systems where 230 ns long. The holo structure at 90 ns was used to model the R241A mutant. The R241A mutant with cAMP was minimized and equilibrated as above and production runs was 200 ns long.

Root mean square deviation (RMSD) was evaluated over the Cα atoms considering all residues between 119 and 370, excluding the six residues of the unstructured N-and C-terminal tails. RMSD based clustering over Cα atoms was performed with Wordom^49^ in the last 50 ns of simulation using a threshold of 2 Å (**Supplementary Fig. 4**).

SMD^50^ simulations were performed starting from a set of conformations from the production phase. For each selected conformation, the protein was centered into the box and rotated to place the Cα atoms of the first and the last residue along the *x*-axis. To take in account the elongation of the protein during the pulling, the box was enlarged, by adding water molecules along the *x* direction, by 170 Å. Finally, the box for SMDs was 260×90×90 Å^3^ and the grid for the PME set to 256×81×81. The solvent was equilibrated for 400 ps harmonically restraining the protein in its original conformation and another 400 ps during which the restraints were progressively turned off. All SMDs were conducted in the NPT ensemble by using the same parameters employed to carried out the MD simulations. The final conformations were then used as starting point for the SMD simulations. The SMDs were performed by restraining the Cα atom of the first residue to its initial coordinates and applying the pulling constant velocity to a dummy atom attached via a virtual spring to the Cα atom of last residue: the spring constant was set to 0.5 kcal mol^-1^ Å^-2^ and the pulling velocity was set to 2.5 Å/ns along the *x* direction (**Supplementary Fig. 4**).

## Acknowledgments

We thank Maria Fe Lanfranco and members of the Maillard and Taylor laboratories for constructive discussions on the manuscript. We thank Amy Chau from the Maillard lab for helping in collecting cAMP titration data for the isolated CNB domains. We thank Emília Pécora de Barros from the Amaro lab at UCSD for providing the atomic coordinates of the simulated R241A structure. We thank Joseph Lesniewski from the Jorabchi lab at Georgetown University and Alan Lowe from University College London for assisting in the global fitting code using PyFolding.

## Author contributions

Y.H. and R.A.M. conceived and designed the research. Y.H. purified and modified all protein constructs. Y.H. and J.P.E performed optical tweezers experiments. L.B. performed the molecular dynamic simulations. Y.H., H.C.H. and R.A.M. analyzed the data. Y.H., E.P., H.C.H., S.S.T. and R.A.M wrote the manuscript.

## Data and materials availability

All data is available in the main text or the supplementary materials.

---

**Supplementary Figure 1.**
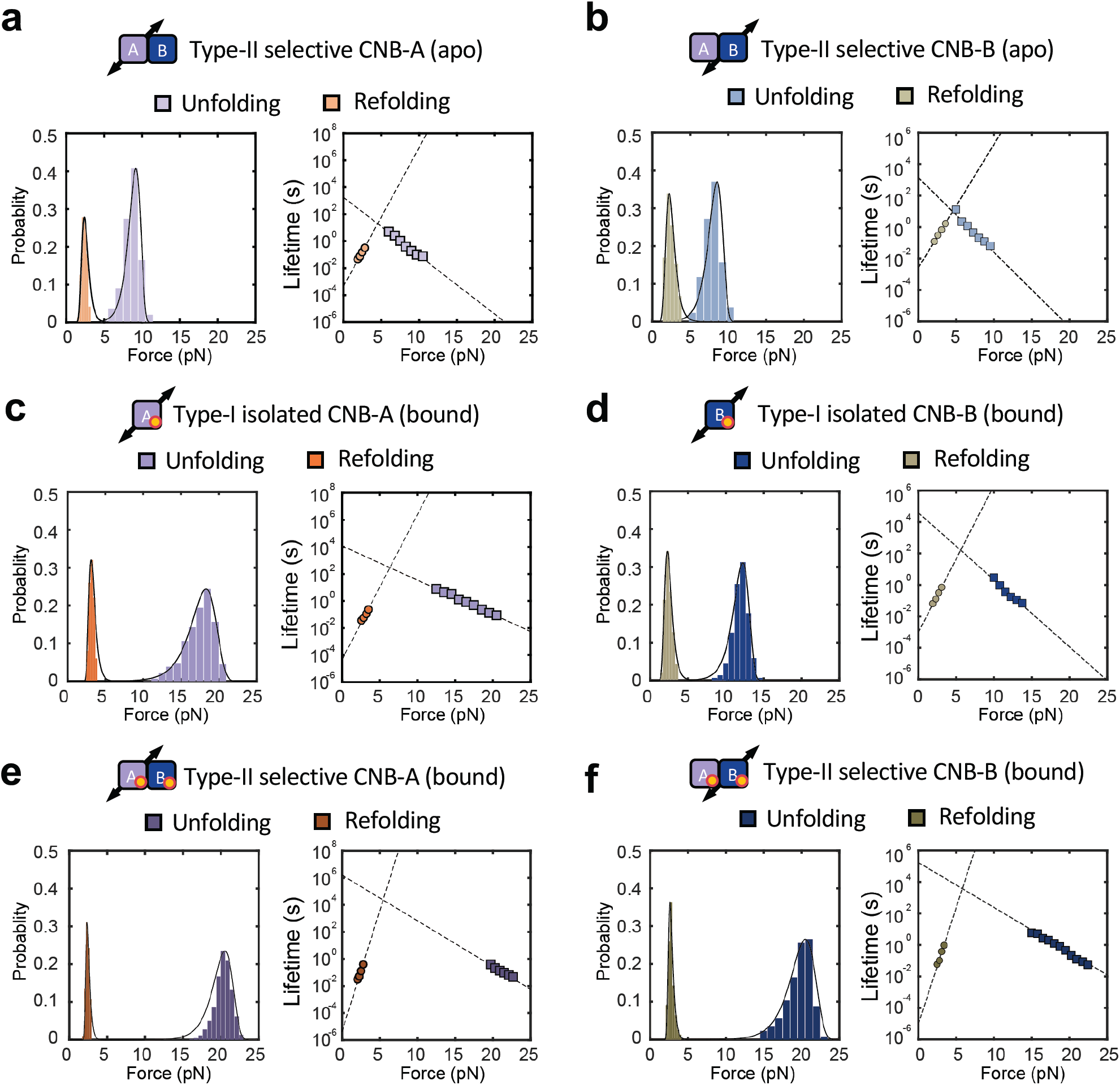
Unfolding and refolding force probability distributions and associated force-dependent folded and unfolded state lifetime with and without cAMP. **a**, CNB-A domain in type-II construct in apo state **b**, CNB-B domain in type-II construct in apo state. **c**, CNB-A domain in type-I construct in cAMP-bound state. **d**, CNB-B domain in type-I construct in cAMP-bound state. **e**, CNB-A domain in type-II construct in cAMP-bound state. **f**, CNB-B domain in type-II construct in cAMP-bound state.

**Supplementary Figure 2.**
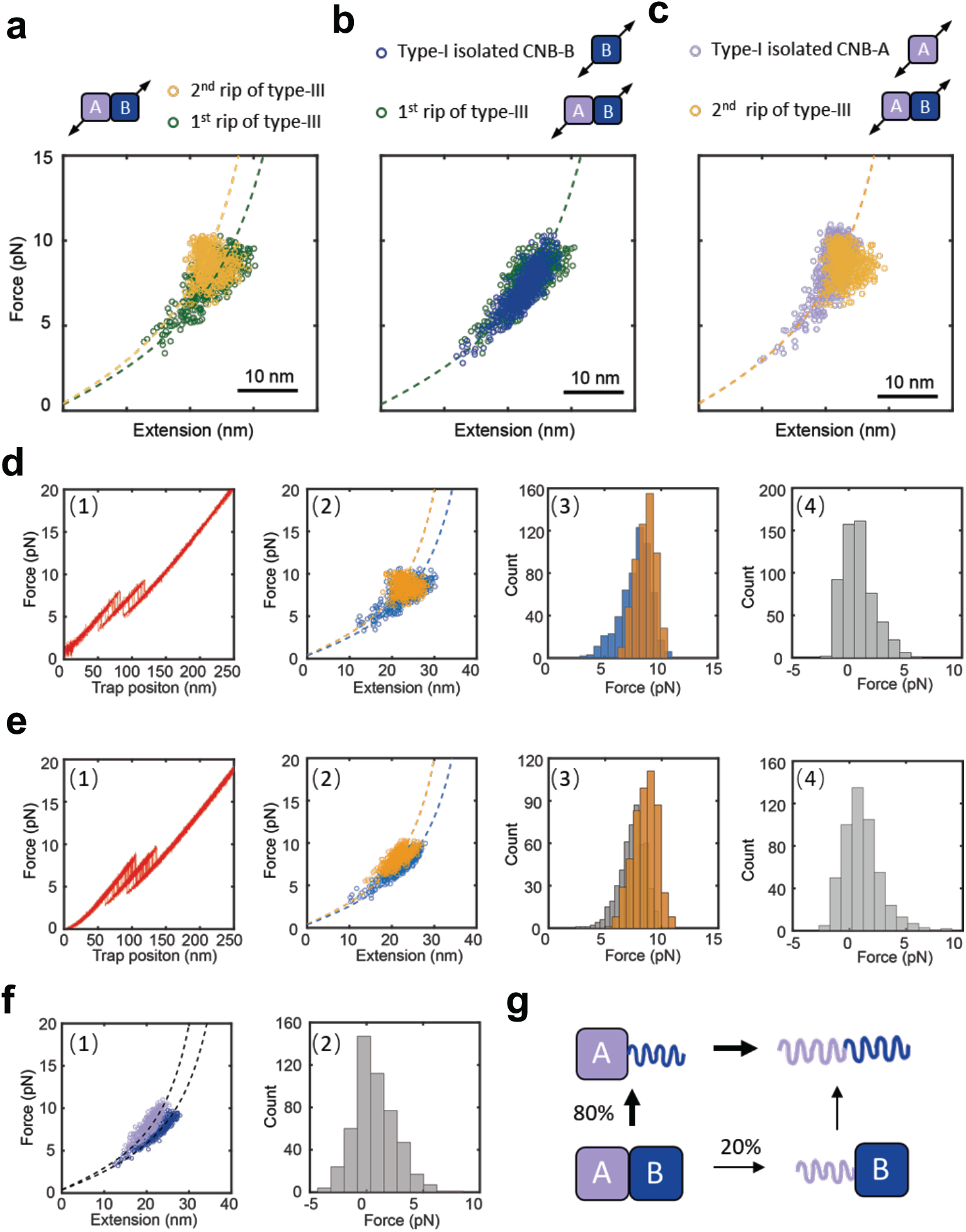
WLC analysis and Monte-Carlo simulations of type-III (S110C/S376C) and type-I constructs. **a**, Change in extension upon unfolding vs. force of the two rips observed in the type-III construct. The dashed lines are the experimentally determined WLC curves for the isolated CNB-A domain (orange, ΔLc = 43 nm) and the CNB-B domain (green, ΔLc = 50 nm). **b**, The 1^st^ unfolding rip in the type-III construct is indistinguishable from those obtained using a type-I CNB-B domain. The dashed line is the experimentally determined WLC curve for the CNB-B domain (green, ΔLc = 50 nm). **c**, The 2^nd^ rip in the type-III construct is indistinguishable from those obtained using the type-I CNB-A domain. The dashed line is the experimentally determined WLC curve for the CNB-A domain (orange, ΔLc = 43 nm). **d**, Experimental data (N=559). (1) Representative force-extension curve obtained with optical tweezers using the type-III construct. (2) WLC analysis of changes in extension upon unfolding vs. force for the 1^st^ (blue) and 2^nd^ rips (orange). The WLC curves were generated using a ΔLc of 43 nm (yellow) and 50 nm (blue) corresponding to the CNB-A and CNB-B domains, respectively (SI Section 7). (3) Unfolding force histogram for the 1^st^ and 2^nd^ rips. (4) Unfolding force difference: 2^nd^ rip minus 1^st^ rip. **e**, Monte-Carlo simulation (N=559). (1) Simulated force-extension curves of the type-III construct were obtained using the kinetic parameters from the individual CNB domains (SI Table 1 and SI Section 8). (2) WLC analysis of the 1^st^ (blue) and 2^nd^ rips (orange) extracted from the simulated force-extension curves. The WLC curves were the same as in 2A. (3) Simulated unfolding force histogram for the 1^st^ and 2^nd^ rips. (4) Simulated unfolding force difference: 2^nd^ rip minus 1^st^ rip. **f**, (1) The Monte-Carlo simulation revealed the identity of the each CNB domain in the plot of changes in extension vs. force. Data for the CNB-B domain is in blue and corresponds to the 1^st^ rip in 80% of all simulated trajectories. Data for the CNB-A domain is in purple. The WLC curves were the same as in 2A. (2) Simulated unfolding force difference between CNB domains: CNB-A minus CNB-B **g**, Cartoon representing the unfolding pathway of the type-III construct reconstructed from the Monte-Carlo simulation. (For detailed analysis, refer to SI Section 2-3)

**Supplementary Figure 3.**
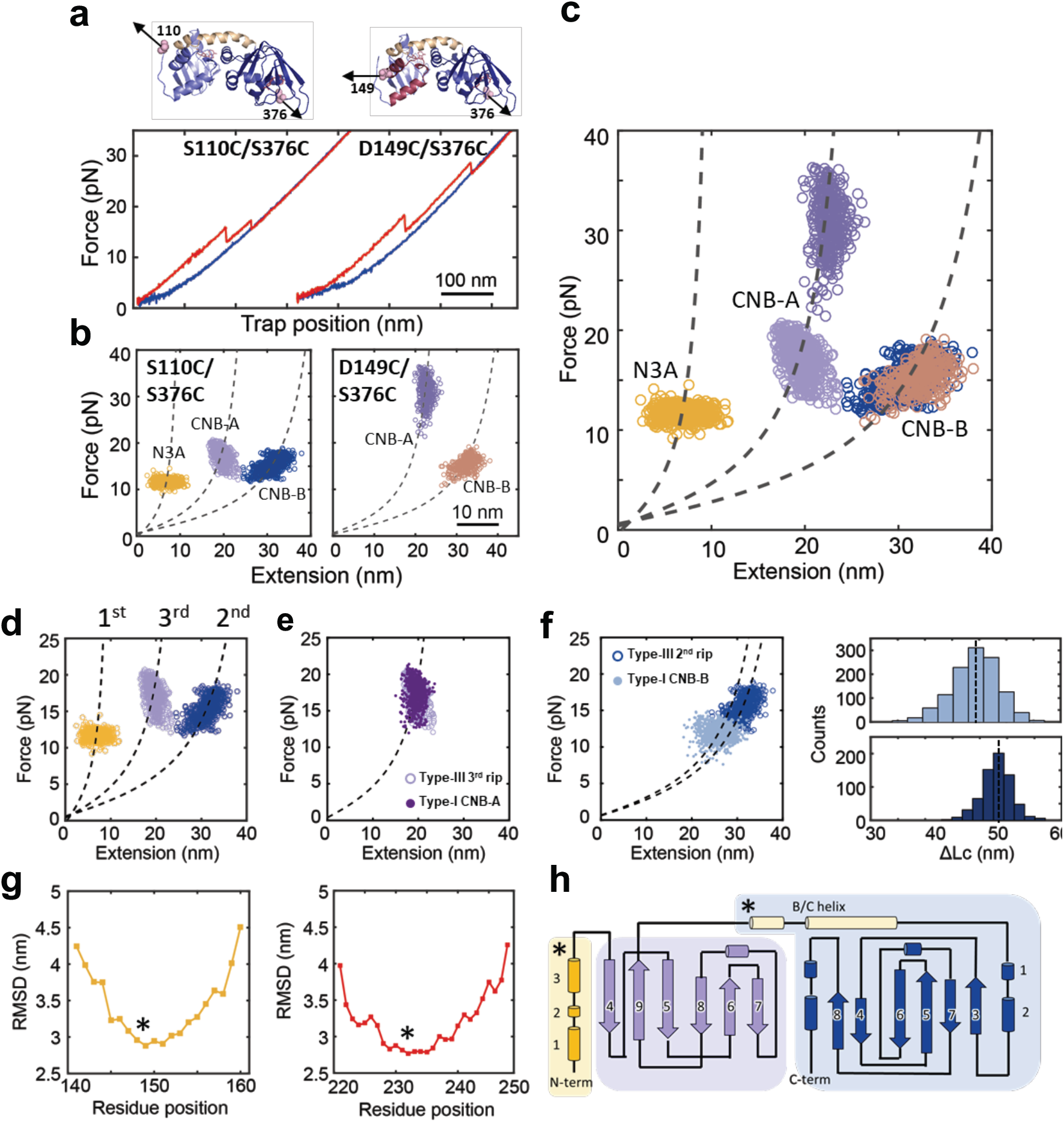
Identification of the N3A motif and assignment of unfolding intermediate for type-III construct in cAMP-bound state. **a**, Force-extension curve of type-III constructs with DNA handles at residue positions S110C/S376C (left) and D149C/S376C (right). Curves were obtained with cAMP. **b**, WLC analysis of changes in extension upon unfolding vs. force obtained for S110C/S376C (left) and D149C/S376C (right). The WLC curves (dashed lines) were obtained using a ΔLc of 13 nm (for the N3A motif), 31 nm (for the CNB-A domain) and 50 nm (for the CNB-B domain). The D149C/S376C does not show a transition corresponding to the N3A motif. **c**, Combined plot of changes in extension upon unfolding vs. force obtained for S110C/S376C and D149C/S376C. **d**, WLC plot of type-III construct (S110C/S376C) with cAMP. The WLC curves (dashed lines) were obtained using a ΔLc of 13 nm (for the N3A motif, yellow), 31 nm (for the CNB-A domain, purple) and 50 nm (for the CNB-B domain, blue). **e**, WLC analysis (using ΔLc = 31 nm) of the changes in extension upon unfolding for the CNB-A domain in the regulatory subunit (type-III construct) overlaid with the data obtained with the isolated CNB-A domain (type-I construct). **f**, WLC analysis (using ΔLc = 47 and 50 nm) of the changes in extension upon unfolding for the CNB-B domain in the regulatory subunit (type-III construct, dark blue symbols) overlaid with the data obtained with the isolated CNB-B domain (type-I construct, light blue symbols). The CNB-B domain in the regulatory subunit has a longer ΔLc (right panels) likely due to the complete folding of the B/C helix. **g**, Mapping the measured ΔLc for each unfolding rip to the corresponding structural elements of the regulatory subunit. Root-mean-square-deviation (RMSD) of the measured extension changes from the predicted WLC extension change as a function of the residue position for the 1^st^ rip (yellow) and 2^nd^ rip (red). The residue that gives the lowest RMSD is labelled with a star. These residues correspond to 149 (1^st^ rip) and 233 (2^nd^ rip). **h**, Topology of the regulatory subunit showing the identified structural elements in each unfolding rip. The 1^st^ rip corresponds to unfolding of the N3A motif (residues 120-149), the 2^nd^ rip corresponds to the CNB-B domain with the BC helix (residues 233-376), and the 3^rd^ rip corresponds to the CNB-A domain minus the N3A motif (residues 150-233).

**Supplementary Figure 4.**
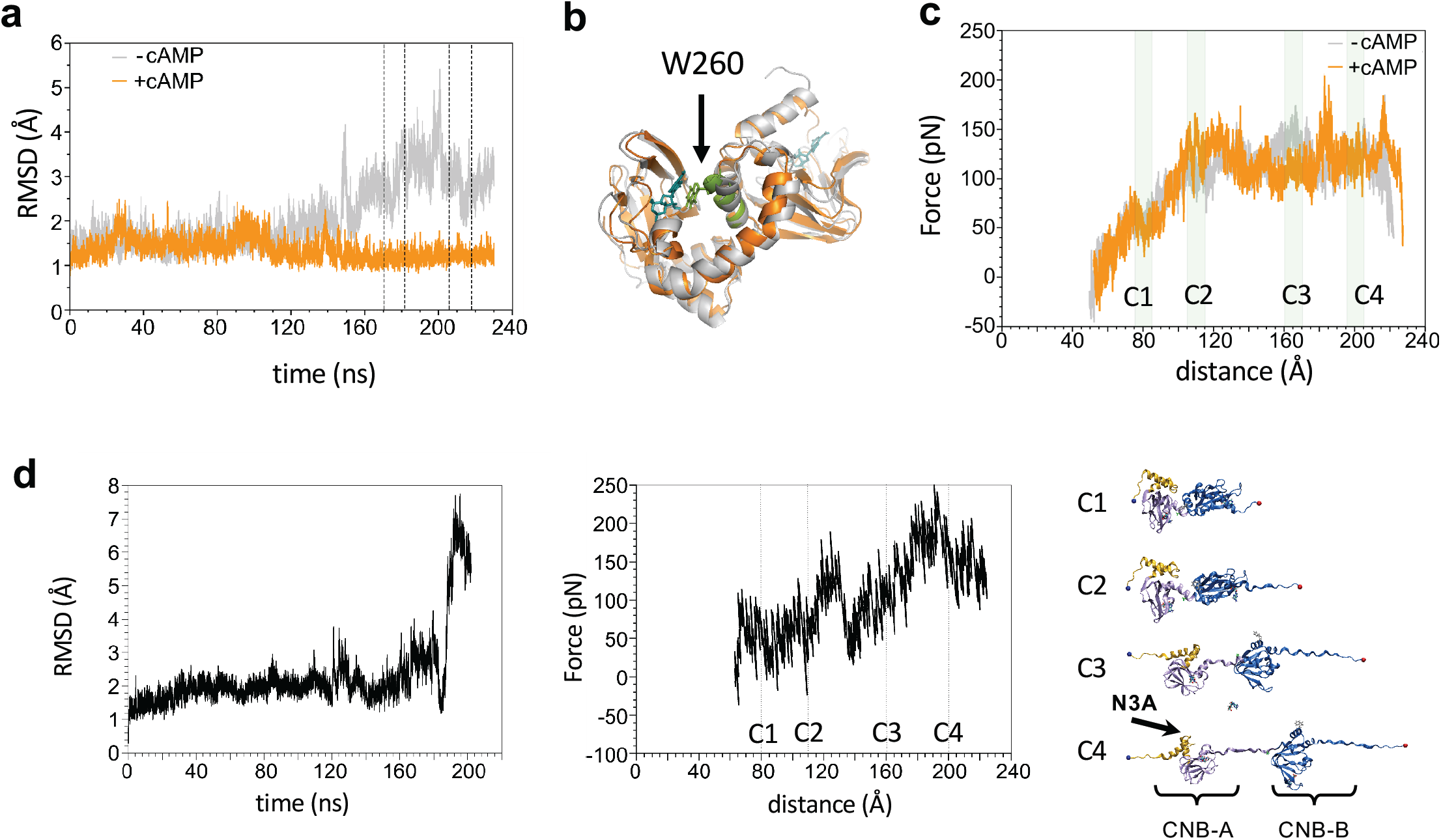
Steered molecular dynamic (SMD) simulations. **a**, Time series of the RMSD from the crystal structure for the PKA regulatory subunit with (orange) and without cAMP (gray). Vertical dotted lines indicate the frames used as starting points for SMD simulations. **b**, Representative structure of the most populated clusters in the last 50 ns of MD simulations with (orange) and without cAMP (gray); highlighted W260 interacting with cAMP docked in the binding site of the CNB-A domain. In the apo state, the lack of the interaction between cAMP and W260 and the absence of cAMP in the CNB-B binding site promote conformational rearrangement of the αA:B helix (green) causing a reciprocal orientation of the domains. **c.** Force-extension profiles for all the SMD simulations with (orange) and without cAMP (gray). Cluster analysis over selected structures (light green shaded areas) were used to characterize the most probable conformations along the trajectories. Yellow: N3A motif; Purple: CNB-A; Dark blue: CNB-B domain. **d**, RMSD time series (left) for MD, force-extension profile for SMD simulations (middle) and selected conformations (gray dashed lines) along the trajectories (right) for mutant R241A with cAMP along SMD.

**Supplementary Figure 5.**
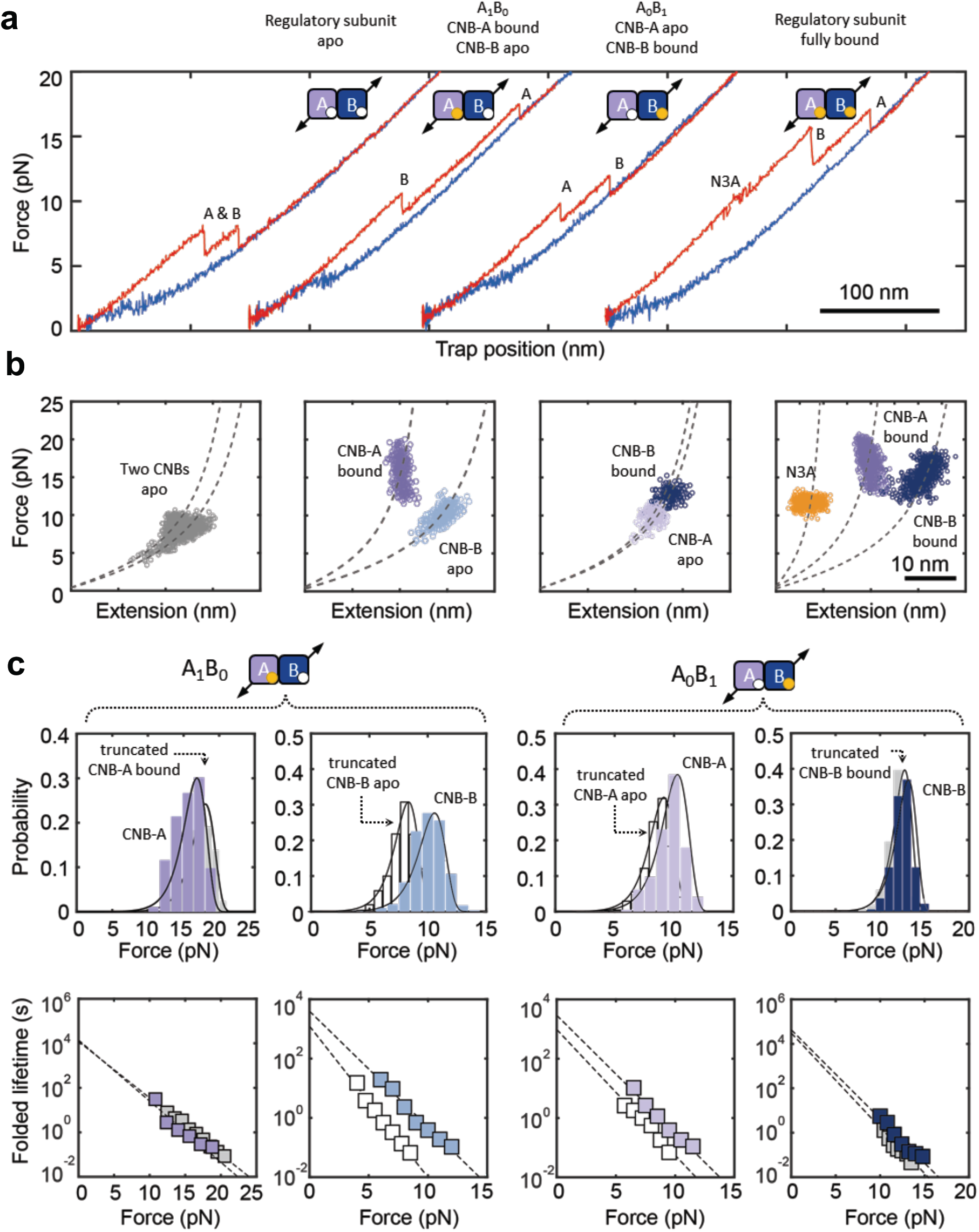
Analysis of intermediate liganded states in cAMP titration of type-III construct. **a**, Representative force-extension curves of four cAMP-bound states using the type-III construct. **b**, Corresponding WLC analysis of changes in extension upon unfolding vs. force. **c**, Unfolding force probability distributions and force-dependent folded state lifetimes of each CNB domain in two intermediate liganded states. The corresponding truncations in the ligand-free (white) and ligand-bound (gray) state were superimposed for comparison.

**Supplementary Figure 6.**
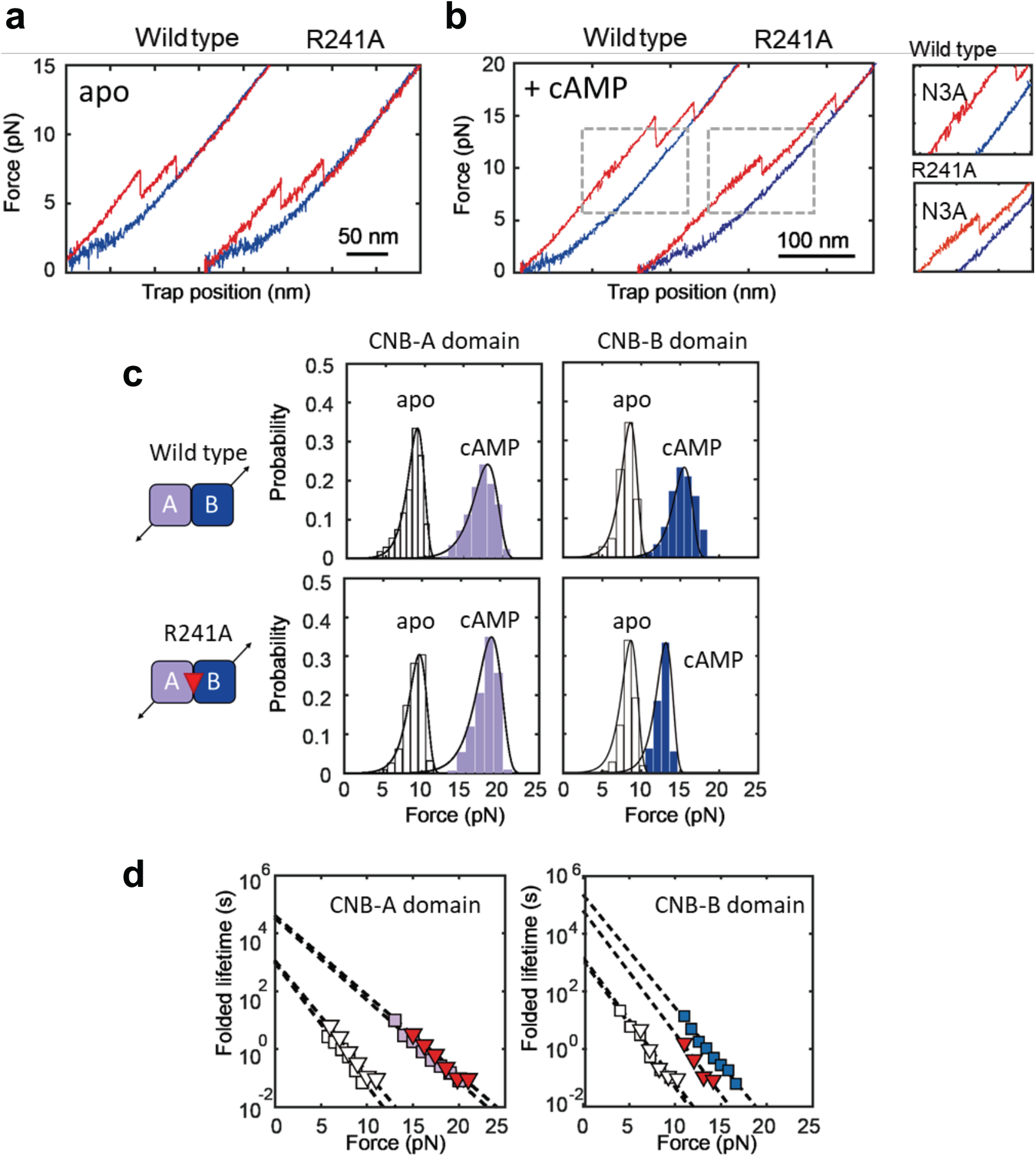
Comparison of type-III (S110C/S376C) wild type and R241A mutant. **a**, Force-extension curves of type-III constructs (S110C/S376C) for wild type and the R241A mutant regulatory subunits in apo state. **b**, Force-extension curves of type-III constructs (S110C/S376C) for wild type and the R241A mutant regulatory subunits obtained with cAMP (left). Zoomed-in trajectories showing the first reversible transition corresponding to the N3A motif in wild type and R241A (right). **c**, Unfolding force probability distributions for each CNB domain corresponding to wild type and R241A. The unfilled bar represents the apo state while filled bars represent cAMP-bound data. **d**, Force-dependent folded state lifetimes for each CNB domain in the apo state (empty symbols) and cAMP-bound conformation (colored symbols) for wild type (squares) and R241A (triangles).

**Supplementary Figure 7.**
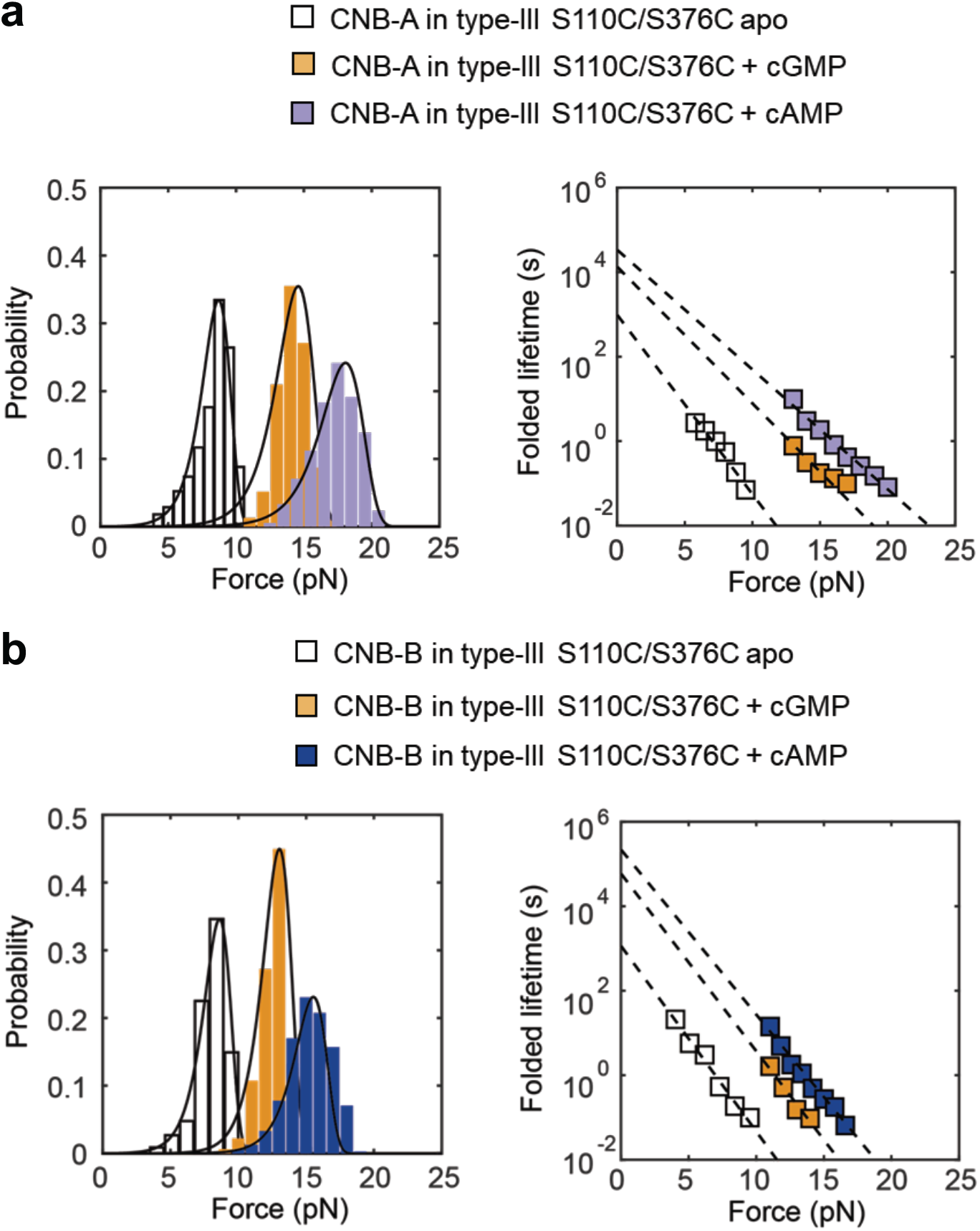
Comparison of unfolding force probability distributions (left) and corresponding force-dependent folded state lifetimes (right) for the type-III construct in apo, cGMP-, and cAMP-bound states. **a**, CNB-A domain and **b**, CNB-B in the regulatory subunit (type-III construct, S110C/S376C) in the apo state (empty symbols) or bound to cGMP (orange) or cAMP (purple and blue).

