## Supplementary Information for "Activation of a Protein Kinase Via Asymmetric Allosteric Coupling of Structurally Conserved Signaling Modules"

#### 1. Quantitative Analysis of Unfolding and Refolding Force Distribution

The force dependent lifetimes were determined from the unfolding force probability histograms using the method from Dudko *et al*[1,2]. Equation (1) is applied considering an unfolding force histogram containing N bins of the width $\Delta F$ that starts at $F_o$ and ends at $F_N = F_0 + N\Delta F$. Let the number of counts in the i$^{th}$ bin be $C_i$, resulting in a height $h_i = C_i/(N_{tot}\Delta F)$ with $N_{tot}$ equal to the total number of counts and where k = 1, 2,…

$$\tau(F_0 + (k - 1/2)\Delta F) = \frac{(h_k/2 + \sum_{i=k+1}^{N} h_i)\Delta F}{h_k\dot{F} + (F_{0\_}(k-1/2)\Delta F} \qquad \text{Equation (1)}$$

where $\dot{F}$ is the force loading rate and $\tau$ is the force dependent lifetime. In order to extract the force dependent lifetimes from the folding force histograms, we used the following equation[3]:

$$\tau(F) = \tau_0\left(1 - \frac{vF\Delta x^{\ddagger}}{\Delta G^{\ddagger}}\right)^{1-1/v} e^{-k_B T\Delta G^{\ddagger}\left[1-\left(1-vF\Delta x^{\ddagger}/\Delta G^{\ddagger}\right)^{1/v}\right]} \qquad \text{Equation (2)}$$

where $\tau_o$ represents the folded lifetime of the protein in the absence of force; F is the force; $\Delta x^{\ddagger}$ is the distance to the transition state from the folded to the unfolded state. $k_B$ is the Boltzmann constant; and T is the absolute temperature. $\Delta G^{\ddagger}$ is the free-energy of activation in the absence of external force. $\nu$ is the scaling factor that specifies the nature of the underlying free-energy landscape. Since the lifetime showed a liner behavior where $\nu$ = 1, the equation (1) was simplified to Bell's model. The unfolding force probability distribution was fitted using the same fitting parameter ($\tau_o$ and $\Delta x^{\ddagger}$) to the following equation[2]:

$$p(\mathrm{F}) = (\mathrm{KV})^{-1} k(F) e^{k_0/\Delta x^{\ddagger} KV} \times e^{-[\frac{k(F)}{\Delta x^{\ddagger} KV}][1-(\frac{vF\Delta x^{\ddagger}}{\Delta G^{\ddagger}})^{1-\frac{1}{v}}]} \qquad \text{Equation (3)}$$

where KV is the force loading rate, $k_0$ is the rate of unfolding at zero force, and k(F) is equal to:

$$k(\mathrm{F}) = k_0 (1 - \frac{vF\Delta x^{\ddagger}}{\Delta G^{\ddagger}})^{\frac{1}{v}-1} \times e^{\Delta G^{\ddagger}[1-\left(1-\frac{vF\Delta x^{\ddagger}}{\Delta G^{\ddagger}}\right)^{\frac{1}{v}}]} \qquad \text{Equation (4)}$$

**2. Worm-like chain (WLC) Analysis for Isolated CNB Domains.**

The extension of the unfolding rip seen in force-extension curves can be analyzed with the Worm-like chain (WLC) model[4] to determine the number of participating amino acids in the unfolding reaction. The WLC equation describes the dependence of force on the molecular extension of a flexible polymer. The resulting force is given by:

$$\mathrm{F} = \frac{k_B T}{p} [\frac{1}{4}\left(1 - \frac{x}{L_C}\right)^{-2} - \frac{1}{4} + \frac{x}{L_C}] \qquad \text{Equation (5)}$$

where $p$ is the persistence length of the chain ($p$ = 0.65nm for polypeptides), $x$ is the end-to-end extension, and $Lc$ is the contour length (calculated by multiplying the number of amino acids by 0.365 nm per amino acid). In calculating changes in contour length:

$$\Delta \mathrm{Lc} = \mathrm{Lc} - d_{folded} \qquad \text{Equation (6)}$$

where $Lc$ is the contour length, and $d_{folded}$ is the end-to-end length between the attachment points in the folded protein (determined from the high-resolution structure).

Therefore, it is possible to compare the number of amino acids that encompass each unfolding reaction under different pulling geometries. By using the WLC model, the truncated CNB-A domain (residues 110-243) has an expected change in contour length of (ΔLc) of 42.8 nm (124 residues × 0.365 nm per residue – 1.89 nm (the distance between residue 110 and 243 in the folded protein)). Since the position 120 is the first structured residue in the CNB-A domain, the total unfolded residue number is 243 – 120 + 1 = 124 instead of 134. In the case of the truncated CNB-B domain (residues 243-376), the expected ΔLc = 48.0nm (134 residues × 0.365 nm per residue – 3.26 nm (the distance between residue 243 and 376 in the folded protein, holoenzyme structure, PDB code: 2QCS[5]).

### 3. Implementation of Monte-Carlo Simulations

The dynamic trajectory of a single tether was carried out by stochastic Monte-Carlo simulations[6,7]. Briefly, simulations were performed by discretization of simulation time into small units $\Delta t$, such that transition probabilities within a given time step were $< 0.05$. For our simulations, $\Delta t$ was chosen to be 5 ms, therefore our simulation was sampled at 200 Hz. Within each time step, iteration of the following processes permitted physical simulation of polymer unfolding:

(1) *Calculation of force-extension for worm-like chains in series*: For each polymer unit in the tension chain (i.e. DNA and protein), a force-extension curve is calculated to relate the polymer unit's fractional extension to applied forces between 0 and 20 pN. At each force, the total extension of the tension chain is the sum of the products of fractional extension and contour length for each polymer unit within the tension chain. Parameters for worm-like chain calculations are provided in a separate paragraph below.

(2) *Stretching of worm-like chains in series*: At each time step, the tension force exerted by the polymer chain is balanced by the pulling force exerted by the optical trap plus a random

fluctuating force ($\vec{F}$): $\vec{F}_{\text{chain}}(t) + \vec{F}_{\text{trap}}(t) + \vec{\xi}(t) = 0$, where *Δ(t)* is a random number chosen from a zero-mean normal distribution with standard deviation $\sigma = \sqrt{2\ k_{\text{B}}T\ \gamma_0/\Delta t}$. The Stokes' drag coefficient, $\gamma_0 = 6\pi r\eta,$ was calculated as a spherical 2.1–$\mu$m diameter bead with radius, $r = 1.05\ \ \mu\text{m}$, in a medium with dynamic viscosity, $\eta = 1\ \ \text{cP}$. The solution to the above force equation was solved numerically at each time step, which also by extension directly calculated the total extension of tension chain and position of the bead in the trap.

(3) *Protein unfolding transition probabilities*: If protein unit A in the tension chain is folded, it is converted to an unfolded state with probability $P(A) = k_A \exp\left[F(t)\Delta x_A^{\ddagger}/k_B T\right]$. Similarly, if protein unit B in the tension chain is folded, it is converted to an unfolded state with probability $P(B) = k_B \exp\left[F(t)\Delta x_B^{\ddagger}/k_B T\right]$.

(4) *Movement of the trapped bead*: The trapped bead is held in an optical trap with Hookean spring constant, $\kappa = 0.075\ \ \text{pN/nm}$. Throughout the simulation, the trap position is moved at a rate of $v = 75\ \ \text{nm/s}$, therefore at each time step, the trap position is incremented by $x(t) = x(t-1) + v\Delta t$.

(5) *Time evolution*: Simulation time *t* was incremented by $\Delta t$.

(6) *Worm-like chain and other parameters used in simulations*. Worm-like chain and parameters used in simulations: 700 bp of DNA was simulated with a persistence length of $P_{DNA}$ = 50 nm, and the unfolded domains with contour lengths L(length $\Delta Lc_{\text{CNB-A}}$) = 46 nm and L($\Delta Lc_{\text{CNB-B}}$) = 52 nm for the CNB-A and CNB-B domains, respectively, had a persistence length of $P_{unf}$ = 0.65 nm. The zero-force protein lifetime for 1,600 s for CNB-A domain, and 1,100 s for the CNB-B domain, leading to zero-force unfolding rates of $k_A = 6.25\times10^{-4}$ $s^{-1}$, and $k_B =$

$9.09 \times 10^{-4}$ $s^{-1}$. The distances to the unfolding transition states used were $\Delta x_A^{\ddagger} = 4.0$ nm, and $\Delta x_B^{\ddagger} = 4.8$ nm. See **Supplementary Table 1** for details.

Discrete time Monte-Carlo simulations were repeated for 2,000 replicates in MATLAB (MathWorks, Natick, Massachusetts), and the features from the resulting stochastic trajectories were plotted directly.

### 4. Assigning the Structures of the Unfolding Intermediates in Type-III Constructs of the Regulatory Subunit Bound to cAMP

We use the WLC model[4] to generate a force versus extension upon unfolding plot from type-III (S110C/S376C) trajectories (**Supplementary Fig. 3**). Three WLC curves were generated using a ΔLc of 13 nm (1st rip), 50 nm (2nd rip) and 31 nm (3rd rip). A similar WLC analysis of the truncated CNB-A and CNB-B domains (type I constructs) bound to cAMP matched the 3rd and 2nd larger rips observed in the type-III S110C/S376C regulatory subunit, respectively. This result indicates that the 2nd rip is due to the unfolding of CNB-B domain while the 3rd rip is originated from the CNB-A domain. Based on the comparison between the type-III S110C/S376C and D149C/S376C constructs (**Supplementary Fig. 3**), the 1st rip corresponds to the N3A motif. The CNB-B domain as part of the regulatory subunit has a longer ΔLc (50 nm) compared to its truncated counterpart (47 nm), likely due to the simultaneous unfolding the CNB-B domain and the B/C helix, which altogether incorporates residues 232 to 376. The distribution of measured contour lengths for the CNB-B domain construct either as a truncation or as part of the regulatory subunit have the expected contour length.

We sought to refine the identity of the secondary structures associated with each unfolding intermediate (or unfolding rip) in the type-III S110C/S376C regulatory subunit. Based on the cAMP-bound high-resolution structure of regulatory subunit (PDB: 1RGS[8]), we mapped the observed rip for each unfolding intermediate to a particular secondary structure using the following equation[9,10]:

$$\Delta L_C = (n+1) \times L_{aa} - (X^D_{m:N\rightarrow C} - X^D_{m+n:N\rightarrow C}) \qquad \text{Equation (7)}$$

$\Delta L_c$ describes the actual change in contour length of each unfolding intermediate (in nm). *n* is the number of residues involved during the transition and $L_{aa}$ is the contour length increment per amino acid (0.365nm/aa). The second term in Equation (7) is the shift in the distance between the last structured residue at the N-terminus (residue position "*m*") and the last structured residue after the unfolding event occurs (residue position "*m*+*n*"). *X* is the folded distance (*D*) from N-terminus to C-terminus (N → C) accounting from residue number *m* or *m*+*n*, which are determined from the crystal structure.

The positions giving the lowest root-mean-square-deviation (RMSD) values from the WLC fit (**Supplementary Fig. 3g, star**) are the optimal residues incorporated in each unfolding intermediate. The 1$^{st}$ rip corresponds to residues 110 to 149, the 2$^{nd}$ rip corresponds to residues 233 to 376, and the 3$^{rd}$ rip corresponds to residues 150 to 232. These assignments corroborate the experimental comparisons among the truncated CNB domains, and the type-III constructs D149C/S376C and S110C/S376C. Each unfolding intermediate was map onto a topological representation of the regulatory subunit topology (**Supplementary Fig. 3h**).

### 5. Contribution of the N3A Motif to the Mechanical Stabilization of the CNB-B Domains

After the N3A motif is fully unraveled, the CNB-B domain unfolds at $F_{avg}$ = 15.0 ± 0.1 pN

(**Fig. 3a, Supplementary Fig. 6c, d** and **Supplementary Table 4**), which is ~ 4 pN lower than the force obtained when the CNB-B domain is selectively unfolded using a type-II construct (**Fig. 2a** and **Supplementary Table 2**). Thus, of the ~ 7 pN of mechanical stabilization conferred by the CNB-A domain, the N3A motif alone contributes 4 pN. The other 3 pN reflect partial stabilization of inter-domain interactions, likely mediated by W260 located in the CNB-B domain that serves as adenine capping residue of the cAMP bound to the CNB-A domain[12,20]. The last step in the unfolding pathway of S110C/S376C corresponds to the CNB-A domain, which occurs at forces similar to those obtained for the isolated domain since other interacting domains are no longer folded (**Fig. 3a** and **Supplementary Fig. 6c, d**).

#### 6. Contact Map Analysis of the PKA Regulatory Subunit

Pairwise contact map comparing the interaction established by the N3A motif in the regulatory subunit for wild type and R241A was analyzed with CMView software[11]. The PDB code used for PKA regulatory subunit is 1RGS. The atomic coordinates of the simulated R241A structure was provided by Emília Pécora de Barros from the Amaro lab at UCSD. The contact map is based on C$\alpha$ atoms using a cutoff distance of 8 Å. The CMView software can be freely obtained at:

http://www.bioinformatics.org/cmview/installation.html

#### 7. cAMP Titration of Type-I and Type-III constructs

Type-I isolated CNB-A and Type-I isolated CNB-B constructs were used in the cAMP titration experiments to determine the binding affinity of each domain. By counting the fraction of bound and unbound at different cAMP concentration, we were able to build a single-molecule titration curve for each isolated domain. The binding affinity of each domain was calculated using the following equation:

$$\text{Fraction Bound} = \frac{k\cdot[cAMP]}{1+k\cdot[cAMP]} \qquad \text{Equation (8)}$$

We obtained $k_{\text{CNB-A}} = 1.2 \pm 0.4\cdot10^7$ M and $k_{\text{CNB-B}} = 3.7 \pm 1.0\cdot10^7$ M for type-I isolated CNB-A and CNB-B respectively (**Supplementary Fig. 5**).

Type-III Regulatory subunit S110C/S376C was used in the cAMP titration experiments (**Fig. 5c**). At different cAMP concentration, every trajectory was assigned to apo ($A_0B_0$), partial liganded state ($A_1B_0$ or $A_0B_1$) and fully bound ($A_1B_1$). Titration curve of each liganded state was plotted, with the error bar showing the weighted STDEV of different molecules.

We established a binding model between cAMP and the two binding sites of the regulatory subunit:

$$\begin{array}{ccc} & k_1 & \\ A_0B_0 + \text{cAMP} & \leftrightarrows & A_{\text{cAMP}}B_0 + \text{cAMP} \\ k_2 \downarrow\uparrow & & \downarrow\uparrow k_3 \\ A_0B_{\text{cAMP}} + \text{cAMP} & \leftrightarrows & A_{\text{cAMP}}B_{\text{cAMP}} \\ & k_4 & \end{array} \qquad \text{(Scheme S1)}$$

Then, each liganded state fraction was determined as a function of cAMP concentration.

$$A_0B_0 = 1/\,(1 + k_1\cdot[\text{cAMP}] + k_2\cdot[\text{cAMP}] + k_1\cdot k_3\cdot[\text{cAMP}]^2)$$

$$A_1B_0 = k_1\cdot[\text{cAMP}]/\,(1 + k_1\cdot[\text{cAMP}] + k_2\cdot[\text{cAMP}] + k_1\cdot k_3\cdot[\text{cAMP}]^2)$$

$$A_0B_1 = k_2\cdot[\text{cAMP}]/\,(1 + k_1\cdot[\text{cAMP}] + k_2\cdot[\text{cAMP}] + k_1\cdot k_3\cdot[\text{cAMP}]^2)$$

$$A_1B_1 = k_1\cdot k_3\cdot[\text{cAMP}]^2/\,(1 + k_1\cdot[\text{cAMP}] + k_2\cdot[\text{cAMP}] + k_1\cdot k_3\cdot[\text{cAMP}]^2)$$

We used *PyFolding* software to proceed the global fitting of four liganded state curves with shared parameters, where $k_1 = 5.7 \pm 0.5 \cdot 10^7$ M$^{-1}$, $k_2 = 1.0 \pm 0.1 \cdot 10^8$ M$^{-1}$, $k_3 = 1.5 \pm 0.2 \cdot 10^8$ M$^{-1}$ and $k_4 = 8.7 \pm 1.2 \cdot 10^7$ M$^{-1}$. The *PyFolding* Script can be freely obtained at: https://github.com/quantumjot/PyFolding[12]

### 8. Bayesian Hidden Markov Model (BHMM) Analysis

We used a Bayesian Hidden Markov Model (BHMM) approach to analyze the optical tweezers data collected under force-clamp experiments[13]. The BHMM analysis method have been previous described and applied in analyzing single molecule trajectories[14]. The MATLAB code used for the BHMM approach can be freely obtained at: http://simtk.org/home/bhmm.

### 9. Calculation of Equilibrium Free Energy

Having obtained the lifetimes of the folded ($\tau_{0,F}$) and unfolded states ($\tau_{0,U}$) at zero force, we estimated the equilibrium free energy of unfolding of each CNB domain using the following equation:

$$\Delta G^0 = -RTln\frac{k_{F\to U}}{k_{U\to F}} = -RTln\frac{\tau_{0,U}}{\tau_{0,F}} \qquad \text{Equation (9)}$$

where RT = 0.592 kcal/mol. The calculations were done with Type-I and Type-II constructs, with or without cAMP. For the CNB-A domain bound to cAMP, $\tau_{0,F}$ was obtained after the N3A motif had unfolded ($\Delta Lc$ = 30.2 nm), whereas $\tau_{0,U}$ was obtained for the full-length protein ($\Delta Lc$ = 43.1 nm). Because these two lifetimes represent different kinetic steps in the unfolding and refolding reactions, and in particular the unfolding reaction does not consider the energy required to unfold the N3A motif, we can only place a lower boundary of $\Delta G^0$ for the cAMP-bound CNB-A domain either as a truncation (Type-I construct) or as part of the PKA regulatory subunit (Type-II construct)(**Supplementary Table 2**). For other conditions (i.e., apo state) or for the CNB-B domain, the folded and unfolded state lifetimes represented the same kinetic step and therefore equation 5 can be directly applied.

**10. Calculation the Change of Solvent Accessible Surface Areas**

The solvent accessible surface areas (ASA) of regulatory subunit bound with cAMP (PDB: 1RGS) and bound with the catalytic subunit (PBD: 2QCS) was calculated. The total change of ASA is the sum of the change in ASA for every residue in the regulatory subunit. The online calculation resources can be freely accessed at: http://curie.utmb.edu/getarea.html[15]

| Region of the change in ASA | Percentage |
|---|---|
| N3A motif | 24% |
| CNB-A domain (not including N3A motif) | 18% |
| B/C helix | 26% |
| CNB-B domain | 33% |

**11. Calculation of the Stability of the N3A Motif in the Regulatory Subunit Bound to cAMP (Type-III S110C/S376C construct)**

Force-clamp data was used to estimate the stability change of the N3A motif from the folded state (F) to the unfolded state (U). This is accomplished using the following the equation[14]:

$$\Delta G_0 = \Delta G_{work} - \Delta G_{stretch} \qquad \text{Equation (10)}$$

where $\Delta G_{work} = F_{1/2} \times \Delta x_{U\text{-}F}$. $F_{1/2}$ is the force in which the folded and unfolded states are equally populated (equilibrium force), and $\Delta x_{U\text{-}F}$ is the extension change difference between U and F. For the wild type regulatory subunit (Type-III S110C/S376C), $F_{1/2}$ = 11 pN with $\Delta x_{U\text{-}F}$ = 5.5 nm, resulting in $\Delta G_{work}$ = 60 pN•nm. Based on the Worm-like chain (WLC) model (SI section 2) and using a persistence length for polypeptides of 0.65 nm, a change in extension of 5.5 nm at 11 pN is equal to a change in contour length of ΔLc = 9.5 nm. $\Delta G_{stretch}$ is the integration of the WLC model using a ΔLc = 9.5 nm with boundaries from F = 0 pN to $F_{1/2}$ = 11 pN, yielding 21.75 pN• nm. Therefore $\Delta G_0$ = (60 – 22) pN•nm = 38 pN•nm = 9.2 $k_BT$ = 5.4 kcal/mol. A similar analysis for the R241A mutation (Type-III S110C/S376C) yielded $\Delta G_0$ = 4.7 kcal/mol.

### Supplementary tables

#### Supplementary Table 1

Kinetic and thermodynamic parameters of type-I (isolated domain) and type-II (selective domain unfolding in PKA regulatory subunit) constructs in the apo state

| Construct | Unfolding Force ± std. (pN) (number of events) | Folded State $\tau_{0,F}$ ± std. (s) | $\Delta x^{‡}_{F\rightarrow U}$ ± std. (nm) | Unfolded State $\tau_{0,U}$ ± std. (s) | $\Delta x^{‡}_{U\rightarrow F}$ ± std. (nm) | $\Delta L_c$± std. (nm) | $\Delta G^0$ (kcal/mol) |
|---|---|---|---|---|---|---|---|
| Type-I: CNB-A | 8.8±1.3 (N=1114) | $1.6{\pm}0.4{\cdot}10^{3}$ | 4.0±0.2 | $2.1{\pm}1.4{\cdot}10^{-4}$ | 9.2±0.9 | 43.1±3.3 | 9.4 |
| Type-I: CNB-B | 7.3±1.3 (N=744) | $1.1{\pm}0.3{\cdot}10^{3}$ | 4.8±0.2 | $3.6{\pm}2.3{\cdot}10^{-3}$ | 6.9±0.9 | 50.3±2.7 | 7.6 |
| Type-II: CNB-A | 8.6±1.2 (N=894) | $2.2{\pm}0.6{\cdot}10^{3}$ | 4.2±0.2 | $4.3{\pm}1.0{\cdot}10^{-4}$ | 9.7±0.3 | 44.3±2.9 | 9.2 |
| Type-II: CNB-B | 7.9±1.1 (N=795) | $1.7{\pm}0.3{\cdot}10^{3}$ | 4.3±0.2 | $2.6{\pm}0.4{\cdot}10^{-3}$ | 7.4±0.2 | 49.9±2.8 | 7.9 |

### Supplementary Table 2

Kinetic and thermodynamic parameters of type-I (isolated domain) and type-II (selective domain unfolding in PKA regulatory subunit) constructs bound to cAMP

| Construct | Unfolding Force ± std. (pN) (number of events) | Folded State $\tau_{0,F}$ ± std. (s) | $\Delta x^{\ddagger}_{F\rightarrow U}$ ± std. (nm) | Unfolded State $\tau_{0,U}$ ± std. (s) | $\Delta x^{\ddagger}_{U\rightarrow F}$ ± std. (nm) | $\Delta L_c$ ± std. (nm) | $\Delta G^0$ (kcal/mol) |
|---|---|---|---|---|---|---|---|
| Type-I: CNB-A | 17.4±2.0 (N=785) | $1.1\pm0.3\cdot10^{4}$ | 2.4±0.1 | $4.4\pm2.4\cdot10^{-5}$ | 10.2±0.6 | 30.2±2.8 | >11.5 |
| Type-I: CNB-B | 12.0±1.0 (N=648) | $3.9\pm0.7\cdot10^{4}$ | 4.0±0.1 | $1.0\pm0.1\cdot10^{-3}$ | 8.8±0.2 | 45.0±2.5 | 10.4 |
| Type-II: CNB-A | 20.3±1.4 (N=1152) | $1.8\pm0.3\cdot10^{6}$ | 3.2±0.2 | $4.1\pm3.0\cdot10^{-6}$ | 16.9±1.1 | 30.7±1.4 | >15.9 |
| Type-II: CNB-B | 19.7±1.6 (N=1518) | $1.4\pm0.5\cdot10^{5}$ | 2.7±0.1 | $1.5\pm1.0\cdot10^{-5}$ | 13.5±0.9 | 47.0±1.9 | 13.6 |

**Supplementary Table 3**

Kinetic parameters of type-III constructs (unfolding both domains simultaneously) in intermediate-liganded states

| Construct* | Unfolding Force ± std. (pN) (number of events) | Folded State $\tau_{0,F}$ ± std. (s) | $\Delta x^{\ddagger}_{F \rightarrow U}$ ± std. (nm) | $\Delta L_c$ ± std. (nm) |
|---|---|---|---|---|
| Type-III: CNB-A ($A_1B_0$) | 15.7±2.0 (N=285) | $1.0 \pm 0.4 \cdot 10^4$ | 2.7±0.3 | 31.3±2.5 |
| Type-III: CNB-B ($A_1B_0$) | 10.0±1.4 (N=285) | $4.0 \pm 2.0 \cdot 10^3$ | 4.6±0.3 | 49.4±2.7 |
| Type-III: CNB-A ($A_0B_1$) | 9.7±1.2 (N=195) | $2.8 \pm 1.9 \cdot 10^3$ | 3.8±0.3 | 44.8±4.8 |
| Type-III: CNB-B ($A_0B_1$) | 12.5±1.0 (N=195) | $5.3 \pm 0.5 \cdot 10^4$ | 3.8±0.1 | 45.5±4.2 |

*Subscripts "1" and "0" denote a domain that is cAMP bound or unbound, respectively.

**Supplementary Table 4**

Kinetic parameters of type-III constructs (unfolding both domains simultaneously) for wild type and R241A bound to cAMP

| Construct | Unfolding Force ± std. (pN) (number of events) | Folded State $\tau_{0,F}$ ± std. (s) | $\Delta x^{‡}_{F \rightarrow U}$ ± std. (nm) | $\Delta L_c$ ± std. (nm) |
|---|---|---|---|---|
| Type-III: CNB-A in wild type | 17.2±2.5 (N=739) | $2.8 \pm 0.5 \cdot 10^4$ | 2.7±0.1 | 30.4±4.7 |
| Type-III: CNB-B in wild type | 15.0±2.2 (N=739) | $1.7 \pm 0.2 \cdot 10^5$ | 3.7±0.1 | 49.4±4.0 |
| Type-III: CNB-A in R241A | 18.2±1.5 (N=339) | $4.0 \pm 0.3 \cdot 10^4$ | 2.6±0.1 | 30.6±1.5 |
| Type-III: CNB-B in R241A | 12.7±0.7 (N=339) | $7.1 \pm 1.3 \cdot 10^4$ | 4.1±0.1 | 49.0±2.4 |

**Supplementary Table 5**

Kinetic parameters of type-I (isolated domains) and type-III (unfolding both domains simultaneously) constructs bound to cGMP

| Construct | Unfolding Force ± std. (pN) (number of events) | Folded State $\tau_{0,F}$ ± std. (s) | $\Delta x^{‡}_{F \rightarrow U}$ ± std. (nm) | $\Delta L_c$ ± std. (nm) |
|---|---|---|---|---|
| Type-I: CNB-A | 12.1±1.1 (N=322) | $7.3 \pm 1.1 \cdot 10^3$ | 3.5±0.1 | 36.8±3.1 |
| Type-I: CNB-B | 10.2±0.8 (N=427) | $3.4 \pm 1.8 \cdot 10^3$ | 3.8±0.2 | 47.5±2.1 |
| Type-III: CNB-A | 14.1±1.1 (N=316) | $1.2 \pm 0.2 \cdot 10^4$ | 3.1±0.1 | 36.1±2.8 |
| Type-III: CNB-B | 12.6±0.9 (N=316) | $6.0 \pm 1.3 \cdot 10^4$ | 4.0±0.1 | 51.7±2.7 |

### SUPPLEMENTARY INFORMATION REFERENCES


1. Dudko, O. K., Hummer, G. & Szabo, A. Theory, analysis, and interpretation of single-molecule force spectroscopy experiments. *Proc. Natl. Acad. Sci.* **105,** 15755–15760 (2008).
2. Dudko, O. K., Hummer, G. & Szabo, A. Intrinsic rates and activation free energies from single-molecule pulling experiments. *Phys. Rev. Lett.* **96,** (2006).
3. Bell, G. I. Models for the specific adhesion of cells to cells. *Science* **200,** 618–627 (1978).
4. Bustamante, C., Marko, J. F., Siggia, E. D. & Smith, S. Entropic elasticity of lambda-phage DNA. *Science* **265,** 1599–1600 (1994).
5. Kim, C., Cheng, C. Y., Saldanha, S. A. & Taylor, S. S. PKA-I Holoenzyme Structure Reveals a Mechanism for cAMP-Dependent Activation. *Cell* **130,** 1032–1043 (2007).
6. Van Kampen, N. G. Stochastic Processes in Physics and Chemistry. *Stochastic Processes in Physics and Chemistry* 96–133 (2007). doi:10.1016/B978-044452965-7/50016-7
7. England, J. P. *et al.* Switching of the folding-energy landscape governs the allosteric activation of protein kinase A. *Proc. Natl. Acad. Sci.* 201802510 (2018). doi:10.1073/pnas.1802510115
8. Su, Y. *et al.* Regulatory subunit of protein kinase A: structure of deletion mutant with cAMP binding domains. *Science* **269,** 807–813 (1995).
9. Sen, M. *et al.* XThe ClpXP protease unfolds substrates using a constant rate of pulling but different gears. *Cell* **155,** (2013).
10. Ries, J., Kaplan, C., Platonova, E., Eghlidi, H. & Ewers, H. A simple, versatile method for GFP-based super-resolution microscopy via nanobodies. *Nat. Methods* **9,** 582–584 (2012).
